# Chromatin Protection by the Chromosomal Passenger Complex

**DOI:** 10.1101/2025.05.15.654082

**Authors:** A. Gireesh, M.A. Abad, R.-S. Nozawa, P. Sotelo-Parrilla, L.C. Dury, M. Likhodeeva, C. Spanos, C. Cardenal Peralta, J. Rappsilber, K.P. Hopfner, M.D. Wilson, W. Vanderlinden, T. Hirota, A.A. Jeyaprakash

## Abstract

The chromosomal passenger complex (CPC; Borealin-Survivin-INCENP-Aurora B kinase) ensures accurate chromosome segregation by orchestrating sister chromatid cohesion, error-correction of kinetochore-microtubule attachments and spindle assembly checkpoint. Correct spatiotemporal regulation of CPC localization is critical for its function. Phosphorylations of Histone H3 Thr3 and Histone H2A Thr120 and modification-independent nucleosome interactions involving Survivin and Borealin contribute to CPC centromere enrichment. However, mechanistic basis for how various nucleosome binding elements collectively contribute to CPC centromere enrichment and whether CPC has any non-catalytic role at centromere remain open questions. Combining a high-resolution cryoEM structure of CPC-bound H3Thr3ph nucleosome with atomic force microscopy and biochemical and cellular assays, we demonstrate that CPC employs multipartite interactions involving both static and dynamic interactions, which facilitate its engagement at nucleosome acidic patch and DNA entry-exit site. Perturbing the CPC-nucleosome interaction compromises protection against MNase digestion in vitro, as well as the dynamic centromere association of CPC and centromeric chromatin stability in cells. Our work provides a mechanistic basis for the previously unexplained non-catalytic role of CPC in maintaining centromeric chromatin critical for kinetochore function.

## Introduction

Faithful distribution of the genetic material between daughter cells during cell division requires correct assembly and regulation of the kinetochore, a multi-subunit protein complex that physically couples chromosomes to the chromosome segregation apparatus, the mitotic spindle (Luykx, 1965; Goldstein, 1981; Uchida *et al*, 2009; Verdaasdonk & Bloom, 2011; Foley & Kapoor, 2013; Fukagawa & Earnshaw, 2014; Musacchio, 2015). The kinetochore is assembled on a specialized chromosomal region, the centromere, which, in most eukaryotes, is specified by the enrichment of nucleosomes containing CENP-A, a histone H3 variant (Earnshaw & Rothfield, 1985; Earnshaw *et al*, 2013; Bodor *et al*, 2014; McKinley & Cheeseman, 2016). In humans and other primates, centromeres are also characterized by the presence of highly repetitive DNA sequences known as α-satellite (Manuelidis, 1978; Fukagawa & Earnshaw, 2014). While the centromere acts as a chromatin platform to assemble the kinetochore, the inner centromere, the chromatin region where the inter-sister chromatid and the inter-kinetochore axis intersect, acts as a key regulatory site where crucial mitotic regulators concentrate (Earnshaw *et al*, 1989; Hindriksen *et al*, 2017). The inner centromere, by recruiting several enzymatic activities (kinases, phosphatases and motor proteins) serves as a signaling platform essential for the regulation of sister chromatid cohesion, kinetochore-microtubule attachments and Spindle Assembly Checkpoint (SAC), processes that are fine-tuned by the interplay between the inner centromere associated kinases and phosphatases (Funabiki & Wynne, 2013; Hengeveld *et al*, 2017). The central component of the inner centromere-associated interaction network is the Chromosomal Passenger Complex (CPC). During prometaphase and metaphase, CPC is enriched at the inner centromere and forms part of a highly regulated interaction network together with Heterochromatin Protein 1 (HP1), cohesin and Shugoshin 1 (Sgo1) (Hindriksen *et al*, 2017).

CPC is composed of two distinct structural and functional modules. The ‘localization module’ includes Survivin, Borealin, and the N-terminal helix of INCENP, which form a triple helical bundle (Klein et al 2006; Jeyaprakash et al., 2007). The ‘kinase module’ comprises Aurora B kinase and the C-terminal region of INCENP, commonly referred to as the INCENP IN-box, which is required for full kinase activation (Bishop & Schuniacher, 2002; Honda *et al*, 2003; Elkins *et al*, 2012).

CPC localization throughout mitosis is highly dynamic, which is directly linked to CPC’s wide ranging functions (Cooke *et al*, 1987; Parra *et al*, 2003). During prophase, CPC localizes along the chromosome arms facilitating chromosome condensation (Lipp *et al*, 2007). During prometaphase and metaphase, CPC is enriched at the inner centromere, where it controls sister chromatid cohesion, destabilizes faulty kinetochore-microtubule attachments (known as error-correction) and regulates SAC activation (Bishop & Schuniacher, 2002; Honda *et al*, 2003; Carmena *et al*, 2012; Hindriksen *et al*, 2017; Haase *et al*, 2017). During late mitosis, CPC translocates to the central spindle and finally accumulates in the midbody during telophase, where it controls cell abscission (Carmena *et al*, 2012; Hadders & Lens, 2022).

Our previous work established that CPC harbors an intrinsic ability to bind nucleosomes through Borealin-mediated multivalent interactions with the histone octamer and the nucleosomal DNA, which are essential for initial chromosome association of the CPC and CPC function (Abad *et al*, 2019). Two mitotic histone marks, phosphorylated Histone H3 Thr3 (H3T3ph) and Histone H2A Thr120 (H2AT120ph), then enrich CPC at the centromere during prometaphase and metaphase. The H3T3ph mark created by Haspin is directly recognized by the BIR domain of Survivin (Kelly *et al*, 2010a; Wang *et al*, 2010; Yamagishi *et al*, 2010; Jeyaprakash *et al*, 2011). The H2AT120ph mark created by the Bub1 kinase recruits Sgo1, which recruits CPC mainly through its interaction with Survivin, with auxiliary contribution from Borealin and INCENP (Klebig *et al*, 2009; Kawashima *et al*, 2010; Tsukahara *et al*, 2010; Yamagishi *et al*, 2010; Trivedi & Stukenberg, 2016; Abad *et al*, 2022). Furthermore, a conserved positively charged ‘RRKKRR’ motif in INCENP (amino acid residues 65 to 70) has also been implicated in centromere enrichment of the CPC (Serena et al, 2020).

Although the enrichment of CPC at centromeres during mitosis and its requirement for accurate chromosome segregation are well established, we still do not understand how various nucleosome-binding elements of CPC subunits cooperatively allow chromatin binding and what CPC’s potential role is in preserving chromatin structure and integrity. Due to the highly repetitive nature of centromeric DNA, centromeres are known to be inherently fragile (Nassar *et al*, 2023). Since protection of centromeric identity is crucial for ensuring accurate chromosome segregation and preserving genome identity, multiple mechanisms are proposed to be at play to ensure the protection and stability of centromeric DNA including DNA methylation, and chromatin binding proteins (Bakhoum & Cantley, 2018; Black & Giunta, 2018; Mellone & Fachinetti, 2021; Nassar *et al*, 2023). Furthermore, considering the microtubule-mediated pulling forces exerted on the centromeric chromatin upon chromosome biorientation, it is likely that inner centromere-enriched mitotic regulators such as CPC, Sgo1 and HP1 contribute to chromatin stability along with cohesin-mediated centromere cohesion (Lera *et al*, 2019; Addis Jones *et al*, 2019). Using structural biology, protein biochemistry and cell biology, here we provide critical insights into the molecular and structural basis for CPC-nucleosome binding and show that stable centromeric association of the CPC is required to stabilize centromeric chromatin, which is crucial for error-free chromosome segregation during cell division.

## Results

### CPC binds nucleosomes through its engagement with the nucleosome acidic patch and the DNA entry-exit site

Several studies, including our own previous work, have established the requirement of the N- terminal tail and the loop region of Borealin (amino acid residues 110-206), the INCENP ‘RRKKRR’ motif (amino acid residues 65 to 70) and the Survivin BIR domain for efficient CPC centromere association (Jeyaprakash *et al*, 2007; Abad *et al*, 2019; Serena *et al*, 2020). However, how the different nucleosome binding elements of CPC collectively contribute towards CPC-nucleosome binding and chromosome association still remains an open question. To address this, we purified recombinant CPC_I1-190SB_ (containing INCENP 1-190, full length Survivin and full length Borealin) which includes the INCENP ‘RRKKRR’ motif and two downstream additional stretches of highly conserved positively charged residues (which we termed basic Intrinsically Disordered Region, basic IDR; Fig. 1A, S1A and S1B), reconstituted a complex with homogeneously modified H3T3 phosphorylated nucleosome core particles (H3T3ph NCPs; using native chemical ligation as described in Abad et al, 2019) (Fig. S1C) and characterized its structure using cryo-Electron Microscopy (cryoEM) (Fig. 1B, S2A, S2B, S2C, S2D, Table 1).

**Fig.1.**
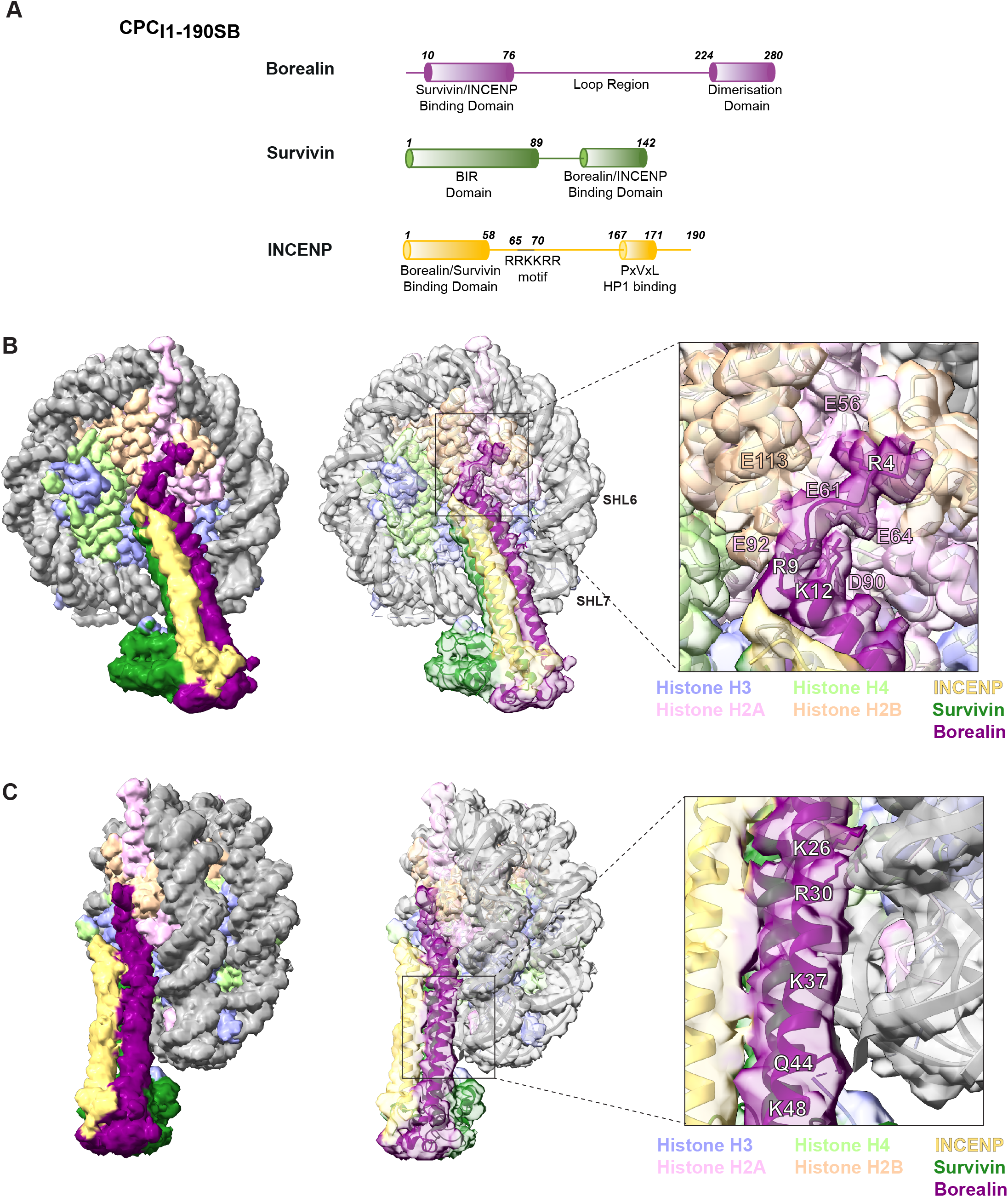
CPC binds to nucleosomes by interacting with both the nucleosome acidic patch and the nucleosome DNA entry-exit site. **(A)** Schematic diagram depicting the domain architecture of the recombinant CPC used in our cryoEM studies, CPC_I1-190SB_, with Borealin full length in purple, Survivin full length in green, and INCENP 1-190 in yellow. **(B)** Top view of the cryo-EM density of CPC_11-190SB_ bound to H3T3ph nucleosome, with the fitted model highlighting the Borealin N -terminal tail and acidic patch interaction. Histones are depicted in violet (Histone H3), light green (Histone H4), pink (Histone H2A), and orange (Histone H2B), and the DNA is shown in grey. CPC subunits are colored in purple (Borealin), green (Survivin), and yellow (INCENP). The zoomed-in image highlights the major residues involved in the salt- bridge interaction, Arg 4, Arg 9 and Lys 12 of the Borealin N-terminal tail (white) making contacts with H2A (pink) and H2B (orange). **(C)** Cryo-EM density (and at a lower contour, and rotated -60 degrees around y-axis with respect to Fig.1B) with the fitted model highlighting the residues on the Borealin helix engaging with DNA between SHL6 and SHL7. The zoomed- in image depicts the side chains of residues, Lys 26, Arg 30, Gln 44 facing the DNA entry-exit site of the nucleosome. Additionally, positions of Lys 37 and Lys 48 are also labelled to emphasize the basic nature of the Borealin region.

**Table 1:** Details of cryo-EM data collection and processing.

|  |  |
| --- | --- |
|  | CPCI1-190SB-H3T3ph NCP Structure |
| Number of grids used | 1 |
| Grid type | Quantifoil R2/2 300 mesh |
| Microscope/detector | Titan Krios / Falcon4i |
| Voltage | 300 kV |
| Magnification | 165k |
| Recording mode | Counting mode |
| Dose rate | 1 e<sup>-</sup>/ Å<sup>2</sup>/frame |
| Defocus | -0.5 μm to -2.6 μm (step size 0.3 μm) |
| Pixel size | 0.727 |
| Total dose | 40 e<sup>-</sup>/Å<sup>2</sup> |
| Number of frames/movie | 40 |
| Total exposure time | Adjusted to keep total dose stable |
| Number of micrographs | 22,065 |
| Number of micrographs used | 12,436 |
| Number of particles used | 73,078 |
| PDB | 9R77 |
| EMDB | EMD-53735 |
| Map resolution (FSC 0.143) | 2.86 |
| **Refinement (Phenix)** |  |
| Resolution (Å) | 3 |
| Map CC | 0.85 |
| Mean B factor (Å<sup>2</sup>) | 98.75 (Protein) 91.57 (Nucleotide) |
| **Validation** |  |
| All atom clashscore | 2.69 |
| Rotamer outliers (%) | 0.11 |
| MolProbity score | 1.23 |
| **Ramachandran plot** |  |
| Favored (%) | 97.04 |
| Outliers (%) | 0.00 |
| **RMS deviation** |  |
| Bond length (Å) | 0.005 (1) |
| Bond angle ( ° ) | 0.703 (0) |

Our 2.8 Å cryo-EM structure revealed that the highly basic Borealin N-terminal tail, with a pI of 12.01, anchors CPC to the nucleosome through its interaction with the acidic patch (Fig. 1B, S2C, S2D, S2E). This mode of anchoring facilitates the engagement of the downstream triple helical bundle formed by Borealin, Survivin and INCENP with the nucleosome DNA entry-exit site (Fig 1C). 3D class analysis of a population of particles with well-defined CPC densities revealed three discrete conformational states of the CPC triple helical bundle (Fig. 2A). This, along with 3D variability analysis, shows that CPC, with the Borealin N-terminal region tethered at the nucleosome acidic patch, swings both vertically and horizontally (Fig. 2A, Movie1). Removing the signal of bound CPC using particle subtraction and subsequent refinement revealed a 2^nd^ copy of the CPC occupying the symmetrically equivalent second face of the nucleosome in a similar conformation (Fig. 2B). In this mode of binding, CPC engages with the DNA entry-exit sites on both sides of the NCP stabilizing the DNA wrapping (Fig. 2B). Interestingly, regions in Borealin and INCENP that were previously identified as essential for CPC binding to nucleosomes (Borealin loop region and INCENP ‘RRKKRR’ motif) were not stabilized in this structure, indicating that they likely form dynamic contacts.

**Fig.2.**
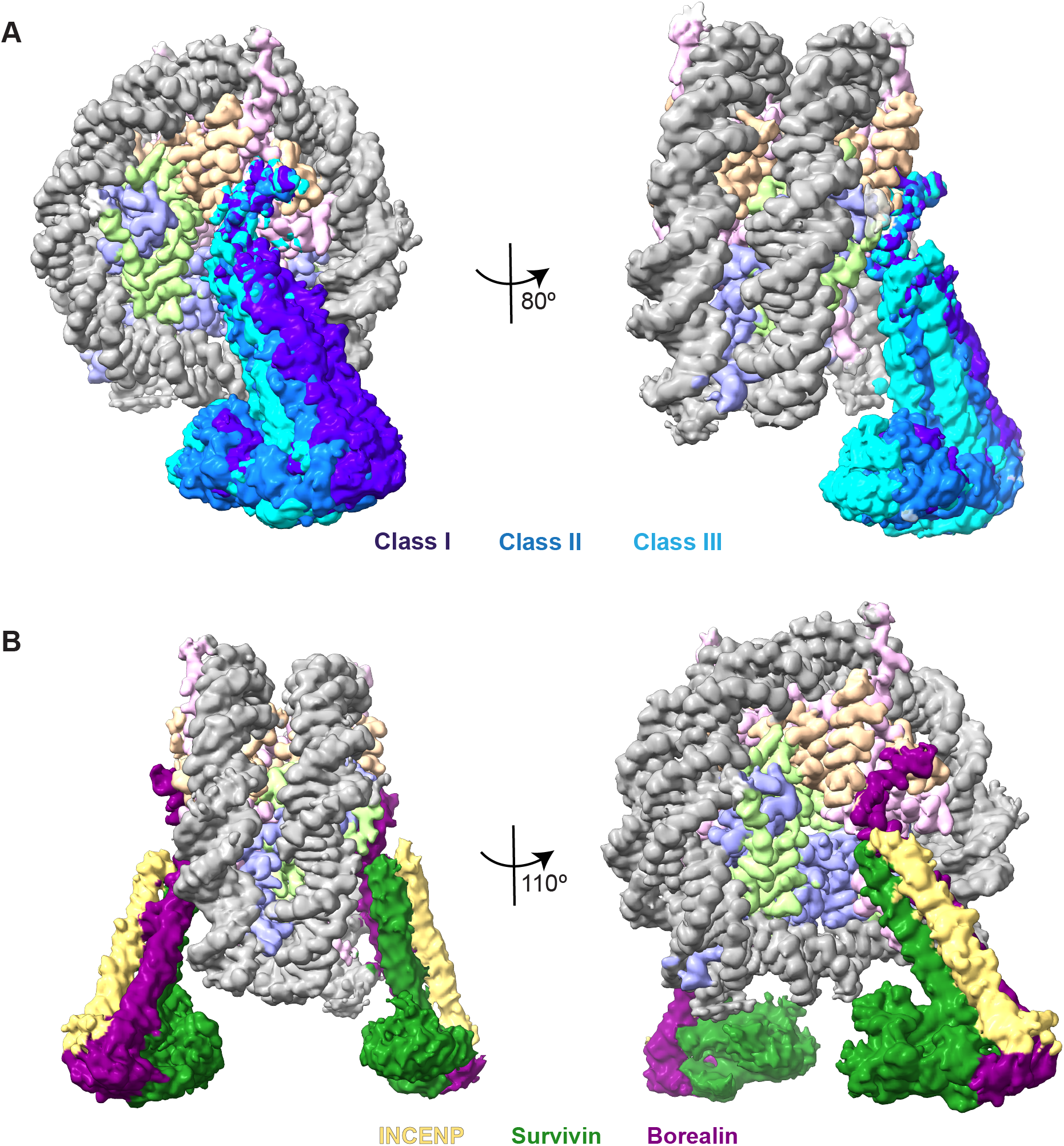
CPC binding is flexible, and CPC occupies both faces of the nucleosome. **(A)** Composite density map of selected classes from 3D classification depicting the swinging motion of CPC. Histones are depicted in violet (Histone H3), light green (Histone H4), pink (Histone H2A), and orange (Histone H2B), and the DNA is shown in grey. The three classes representing the separate positions of CPC are shown in different shades of blue. CPC has flexibility in both directions, both horizontally and vertically, while being tethered to the nucleosome via the N-terminal tail of Borealin. The left image depicts the horizontal flexibility of the complex, while the right image shows the vertical motion of CPC. **(B)** Composite density map depicting the double occupancy of CPC. CPC densities can be seen binding to both faces of the nucleosome. The nucleosome components and CPC densities are colored as in Fig. 2B.

A detailed analysis of the intermolecular interactions at the NCP acidic patch showed that Borealin N-terminal tail amino acid residues Arg 4 and Arg 9 form the arginine anchor that interacts with the H2A/H2B Asp/Glu residues of the nucleosome acidic patch, through salt bridge interactions (Fig. 1B). Borealin Arg 4 interacts with Glu 113 of H2B, and Glu 56 of H2A, while Borealin Arg 9 contacts with Glu 61, Asp 90 and Glu 92 of H2A (Fig. 1B). In addition to the arginine anchor, K12 of Borealin interacts with Glu 61 and Glu 64 of H2A (Fig. 1.B). The CPC_I1-190SB_-H3T3ph NCP structure also revealed that the Borealin helix facing the DNA entry-exit site is highly basic with residues Lys 26, Arg 30, Lys 37 and Lys 48, among which Arg 30 makes Van der Waals contacts with the phosphate backbone of the DNA between super-helical location (SHL) 6 and 7 (Fig. 1C). Additionally, Gln 44 of Borealin makes H-bonding interactions with the backbone phosphates of nucleotide at the DNA entry-exit site (Fig. 1C).

### Dynamic interactions are required for efficient CPC-nucleosome binding

Considering that the anchoring of CPC at the NCP acidic patch is mediated via a well-defined network of electrostatic interactions involving Borealin N-terminal tail residues, we wondered if the Borealin-NCP acidic patch interactions are sufficient to achieve CPC-NCP binding. To test this, we reconstituted nucleosomes harboring acidic patch mutations (Glu 56, Glu 61, Asp 90 and Glu 91 on H2A, and Glu 105 and Glu 113 on H2B, all mutated to Ala) and studied the effect on CPC binding to nucleosomes using Electrophoretic Mobility Shift Assays (EMSA) (Fig. 3A). Interestingly, the acidic patch mutations did not abolish CPC binding to nucleosomes, suggesting that other CPC regions also contribute to NCP binding. Furthermore, we purified a CPC complex lacking the N-terminal 10 residues of Borealin (CPC_I1-190SB10-end_) and tested its ability to interact with H3T3ph NCPs using size exclusion chromatography (SEC) (Fig. 3B). SEC analysis shows that CPC_I1-190SB10-end_ can form a complex with H3T3ph NCPs (Fig. 3B). This observation suggests that multivalent and dynamic interactions involving different regions of Borealin and INCENP, not involving the nucleosome acidic patch, are essential for CPC-NCP binding (Abad *et al*, 2019; Serena *et al*, 2020). These dynamic interactions, likely involving protein-protein and protein-DNA contacts, may facilitate high-affinity binding of CPC to nucleosomes.

**Fig. 3.**
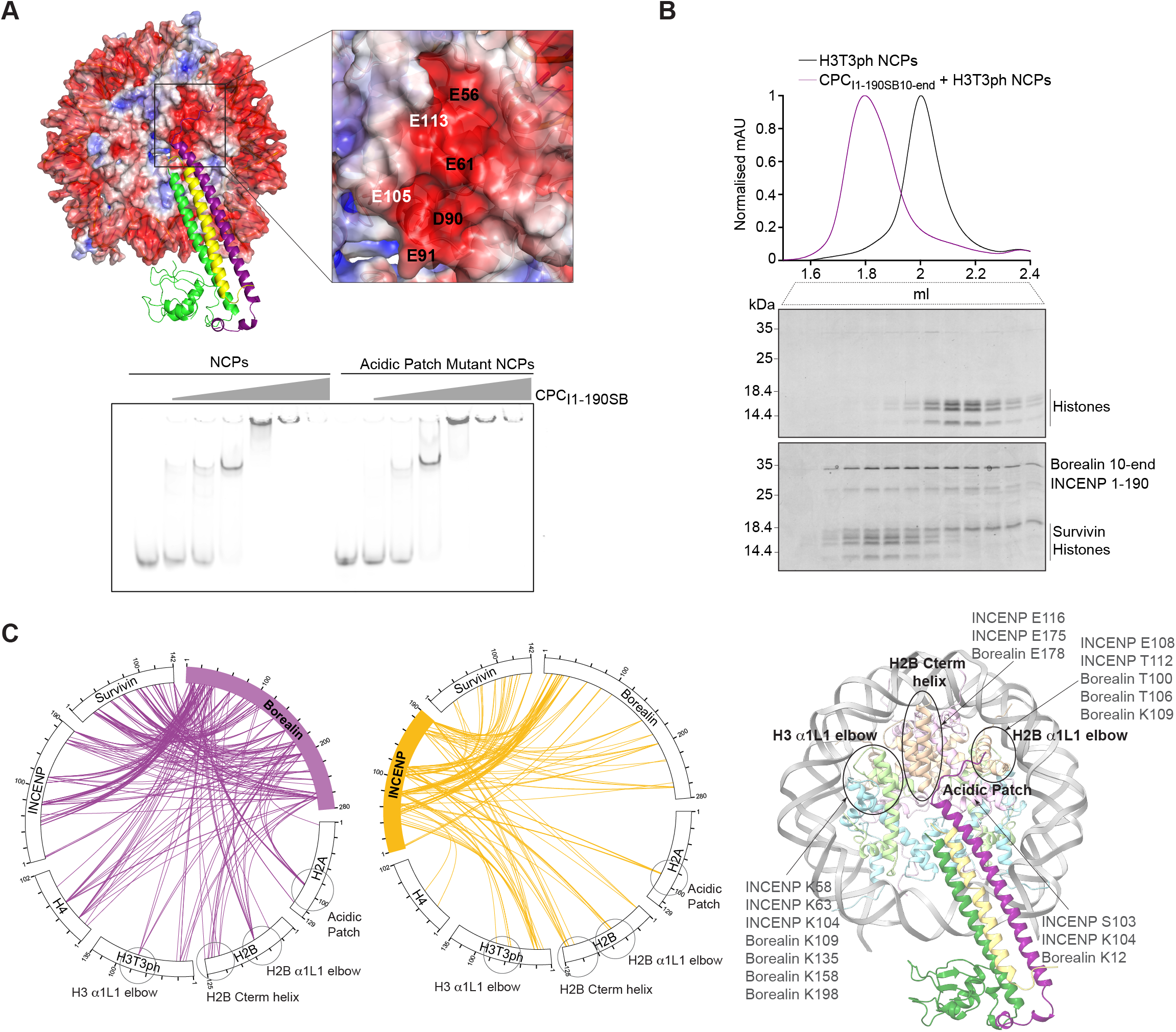
Static nucleosome interactions are not the sole contributors towards CPC-NCP binding. **(A)** Representative native PAGE of EMSA assay performed with 20 nM wild type NCPs and Acidic patch mutant NCPs (Glu 56, Glu 61, Asp 90 and Glu 91 on H2A, and Glu 105 and Glu 113 on H2B all mutated to Ala) with increasing amounts of recombinant CPC_I1-190SB_ (from 20 nM to 640 nM). Top panel shows the Cryo-EM model surface colored by electrostatic potential (APBS using Pymol version 3.1) and showing the acidic patch residues mutated in the EMSA analysis (H2A in black and H2B in white). **B)** Size Exclusion Chromatogram and corresponding SDS-PAGE analysis for H3T3ph nucleosomes (black) and CPC_I1-190SB10-end_-H3T3ph NCP complex (purple). The shift on the purple profile towards the left indicates complex formation, as observed in the SDS-PAGE. **(C)** Circular diagram of the crosslinks observed for the CPC_I1-190SB_-H3T3ph NCP complex. Borealin and INCENP crosslinks with histones are shown in purple and yellow, respectively. On the right, crosslinked regions between CPC components and the histones have been mapped on the cryo-EM model. The four main hotspots of nucleosome binding are highlighted with circles on the crosslinking diagram and the model: the acidic patch, the Histone H3 α1L1 elbow, the Histone H2B C-term helix, and the Histone H2B α1L1 elbow. Both Borealin and INCENP make extensive contacts with these hotspots.

To assess the contribution of dynamic interactions crucial for CPC-nucleosome binding, we performed crosslinking/MS experiments with the CPC_I1-190SB_-H3T3ph NCP complex (Fig. 3C and Fig. S3A). The crosslinking/MS data demonstrated that the additional CPC contacts, involving the Borealin loop region and the INCENP basic IDR, include nucleosome regions previously identified as hotspots for NCP-binding proteins, the H2B C-terminal helix, the H2B α1L1 elbow and the H3 α1L1 elbow (McGinty & Tan, 2021) (Fig. 3C). These nucleosome hotspots mediate intermolecular interactions crucial for various cellular processes, including gene regulation, chromatin remodeling and DNA repair (Lobbia *et al*, 2021).

We have previously shown that Borealin can directly bind to DNA (Abad *et al*, 2019). In addition, the INCENP ‘RRKKRR’ motif has also been suggested to mediate DNA binding (Serena *et al*, 2020). UV-crosslinking of CPC-NCP complex, followed by SDS-PAGE analysis of the nucleosomal DNA revealed the presence of DNA crosslinked with Borealin migrating at the expected molecular weight (Fig. 4A and S3B). The same analysis with CPC lacking the Borealin loop region, resulted in a crosslinked DNA species with faster migration in proportion with the reduced molecular weight of Borealin lacking the loop region (Fig. 4A). Consistent with this, the MS analysis of the UV crosslinked CPC-NCP sample showed a significant reduction in the amounts of unmodified Borealin peptides (i.e., non-DNA-bound peptides) compared to INCENP peptides, likely due to an increase in Borealin peptides containing modification(s) (peptides crosslinked to DNA) (Fig. 4A and S3B), while INCENP levels (based always on unmodified peptides) were largely unchanged. This suggests that INCENP contributes to CPC-nucleosome binding mainly through histone contacts, while Borealin contributes through both DNA and histone contacts.

**Fig.4.**
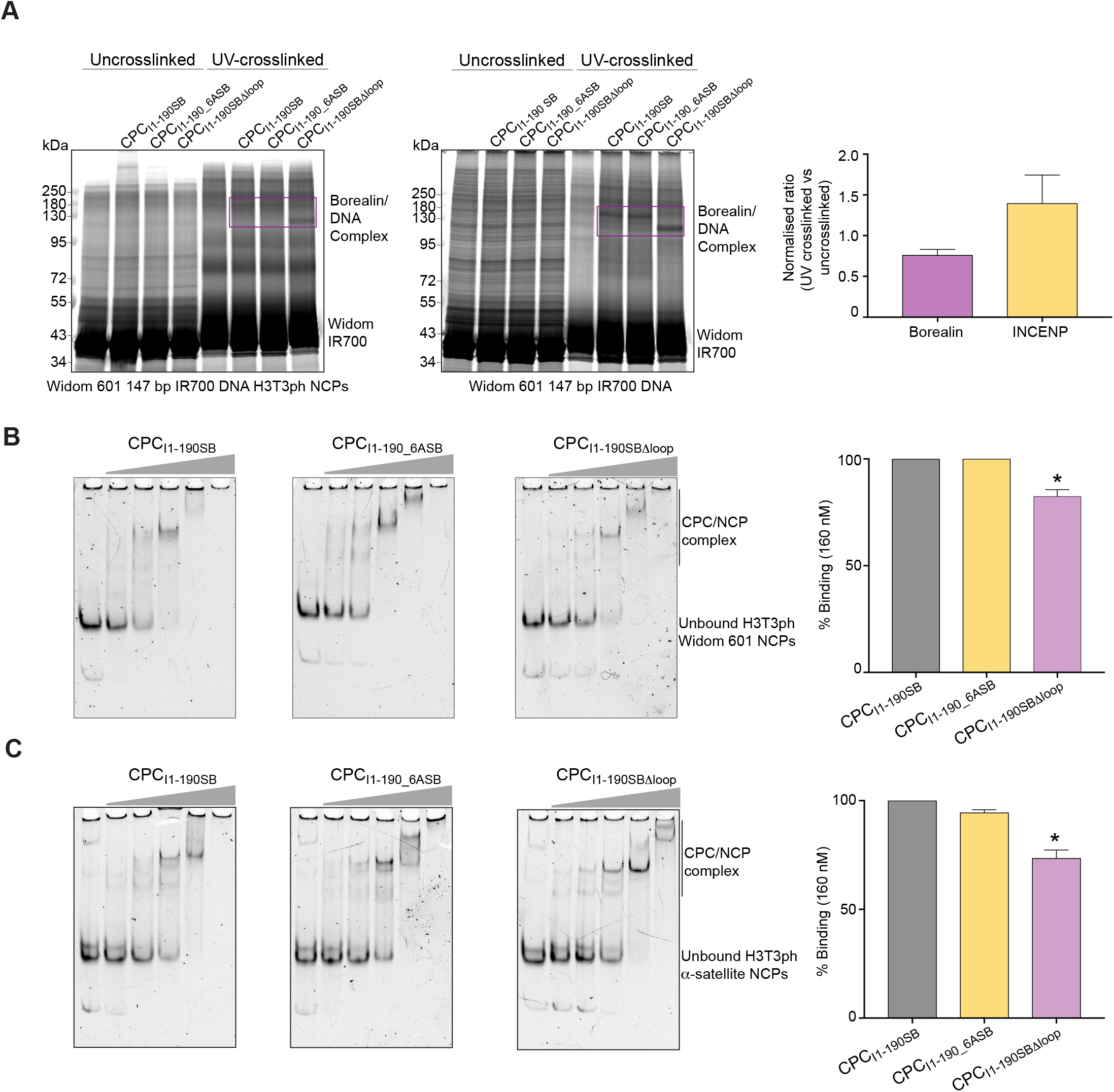
Efficient CPC-nucleosome binding relies on dynamic interactions. **(A)** Representative SDS-PAGE gels of UV-crosslinking analysis of CPC complexes (CPC_I1-190SB_, CPC_I1-190_6ASB_, and CPC_I1-190SBΔloop_) with IR700-labelled 601 Widom 147 bp H3T3ph NCPs (left) or with IR700-labelled Widom 601 147 bp DNA (right). The two Borealin-DNA crosslinked bands appearing at around 130 kDa (for CPC_I1-190SB_ and CPC_I1-190_6ASB_ samples) and 110 kDa (for the CPC_I1-190SBΔloop_ sample) are highlighted inside the box. Molecular weight of Borealin and Borealin 1′loop are 31.32 kDa and 20.57 kDa, respectively. The molecular weight for dsDNA Widom 601 is 91.42 kDa. Corresponding MS/MS quantification of the UV-crosslinked/not UV-crosslinked ratio of Borealin (purple) and INCENP (yellow) protein intensities normalized to the histones within the sample is shown on the right. **(B)** Representative native gels of EMSAs performed with H3T3ph NCPs wrapped with Widom 601 (top) and centromeric α-satellite (bottom) DNA in the presence of increasing amounts of three different CPC constructs (CPC_I1-190SB_, CPC_I1-190_6ASB_, and CPC_I1-190SBΔloop_). Percentage of nucleosome binding at 160 nM CPC concentration is quantified on the right. Kruskal–Wallis with Dunn’s multiple comparisons test; *, P ≤ 0.05. (**C**) Representative native gels of EMSAs performed with H3T3ph NCPs wrapped with α-satellite DNA in the presence of increasing amounts of three different CPC constructs (CPC_I1-190SB_, CPC_I1-190_6ASB_, and CPC_I1-190SBΔloop_). Percentage of nucleosome binding at 160nM CPC concentration is quantified on the right. Kruskal–Wallis with Dunn’s multiple comparisons test; *, P ≤ 0.05.

To directly assess the dynamic contribution of INCENP ‘RRKKRR’ and the Borealin loop region for nucleosome binding, we performed EMSA assays with different versions of CPC containing either mutations or deletions of key INCENP and Borealin regions (CPC_I1-190/6ASB_ and CPC_I1-190SBΔloop_) and H3T3ph nucleosomes reconstituted with either Widom 601 or centromeric α-satellite DNA (Fig. 4B, Fig. 4C, and Fig. S3C and S3D). EMSA analysis showed that while CPC_I1-190/6ASB_ bound to NCPs with a similar affinity to the wild type CPC complex (CPC_I1-190SB_), the CPC construct lacking the loop region of Borealin (CPC_I1-190SBΔloop_) showed a 3-fold decrease in affinity compared to CPC_I1-190SB_. These observations suggest that while Borealin loop region contributes to CPC-nucleosome affinity, the dynamic contacts involving INCENP ‘RRKKRR’ might contribute to the dynamics of CPC-NCP binding.

Altogether, our structural and biochemical analysis shows that CPC-NCP binding is mediated by multi-partite interactions involving both stable and dynamic interactions. This, along with our observation that CPC engages with the nucleosome DNA entry-exit site on both faces of the NCP suggests that CPC likely stabilizes chromatin by protecting the wrapped state of nucleosomal DNA.

### CPC stabilizes nucleosomes through compaction

Atomic Force Microscopy (AFM) enables the visualization of DNA wrapping states around histone proteins within asymmetrically wrapped nucleosomes, providing insights into their conformational landscape (Lyubchenko, 2014; Konrad *et al*, 2021a, 2022). To assess if CPC binding stabilizes nucleosomal DNA wrapping, we performed AFM imaging experiments with H3T3ph NCPs reconstituted with a 486 bp DNA construct (comprising the Widom 601 nucleosome positioning sequence flanked by a short-106 bp and long-233 bp extra-nucleosomal DNA arm; Fig. 5A) in the absence and presence of CPC_I1-190SB_. In the absence of CPC_I1-190SB_, we found that 40 ± 1% (error is SD) of H3T3ph nucleosomes occur in the fully wrapped state, and 60 ± 1% occur in a partially unwrapped state. Addition of CPC_I1-190SB_ increased the fraction of fully wrapped H3T3ph nucleosomes to 52 ± 2% (Fig. 5B, 5C and Fig. S4A-E). Moreover, the presence of CPC_I1-190SB_ led to a highly significant change in the nucleosome wrapping landscape towards a smaller opening angle (Fig. 5B, 5D and Fig. S4A-E), strengthening our hypothesis that CPC binding stabilizes the fully wrapped state of nucleosomes.

**Fig. 5.**
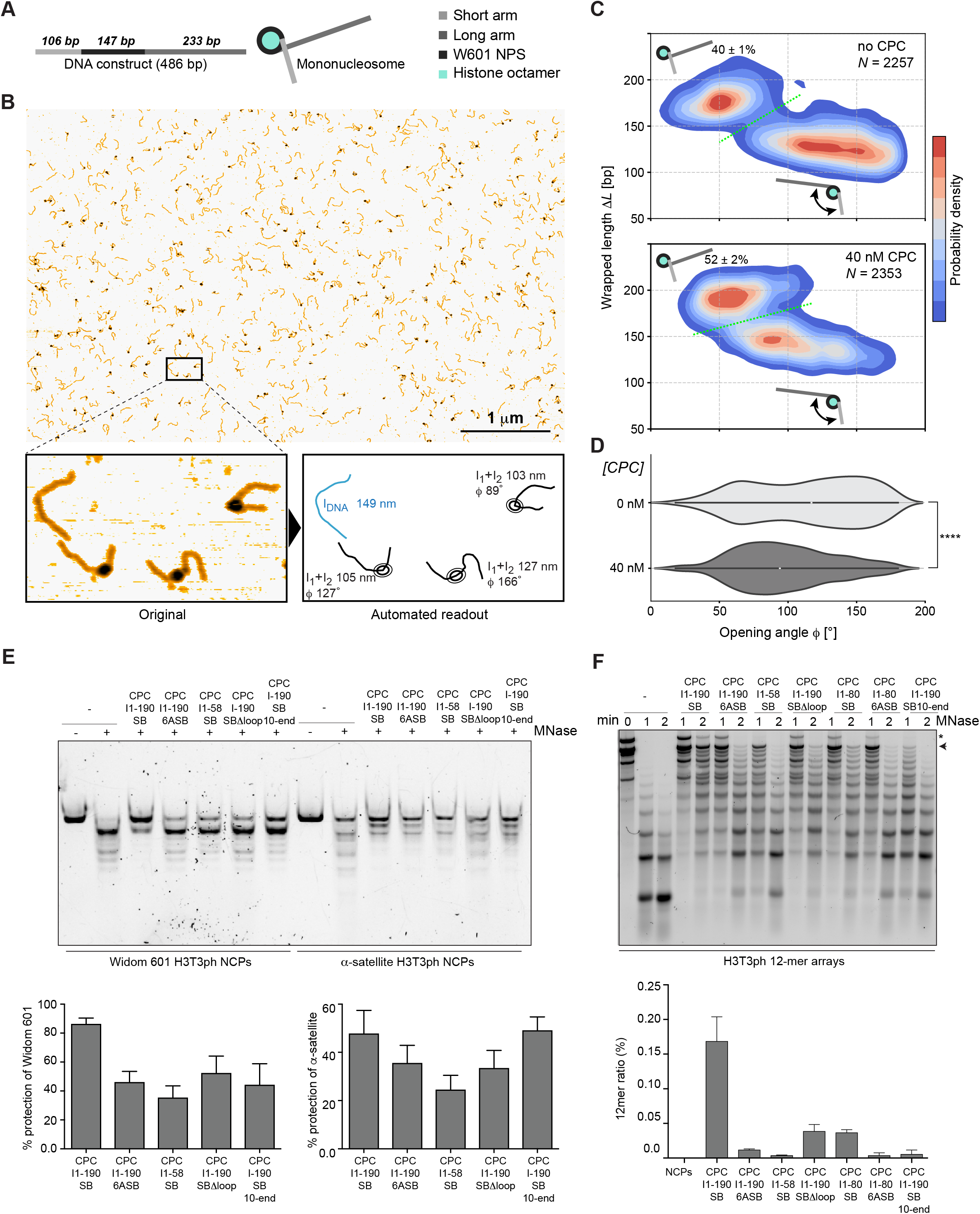
CPC stabilizes highly wrapped states of H3T3ph nucleosomes. **(A)** Schematic depiction of the DNA construct used for nucleosome reconstitution for high-throughput AFM studies. **(B)** Representative overview image of H3T3ph nucleosomes incubated with 40 nM CPC_I1-190SB_. Digital zoom of the boxed region is shown below, depicting three mono-nucleosomes and a bare DNA molecule, before and after automated read-out. **(C)** Two-dimensional kernel density estimates of wrapped length ΔL = <L_DNA_>-l_1_-l_2_ versus opening angle of H3T3ph nucleosomes in the absence (top) and presence of 40 nM CPC_I1-190SB_. The green dotted line indicates the cut-off separating the fully wrapped and partially unwrapped nucleosomes. **(D)** Violin plots of nucleosome opening angles in the absence and presence of CPC_I1-190SB_. A two-sided Kolmogorov-Smirnov statistical test demonstrates a significant difference (p = 5×10^-49^). **(E)** Native gel image of the MNase assay of H3T3ph mono-nucleosomes with Widom 601 147bp DNA and α-satellite DNA in the presence of five different CPC constructs (CPC_I1-190SB_, CPC_I1-190_6ASB_, CPC_I1-58 SB_, CPC_1-190SBΔloop_, and CPC_1-190SB10-end_). Bottom panel, bar graph with the respective quantification of the percentages of DNA protection observed. **(F)** Representative agarose gel image of the MNase assay with H3T3ph 12-mer nucleosomal arrays in the presence of different CPC constructs: CPC_I1-190SB_, CPC_I1-190_6ASB_, CPC_I1-58SB_, CPC_1-190SBΔloop,_ CPC_I1-80SB_ (lacking the basic IDR regions of INCENP downstream of RRKKRR motif), CPC_I1-80_6ASB_ (lacking all positive stretches in INCENP), and CPC_1-190SB10-end_. The corresponding bar graph with the quantification of protection of the 12-mer band (highlighted with the arrow in the gel) is shown below.

We then assessed the extent of nucleosomal DNA protection conferred by CPC binding. We incubated CPC complexes containing different INCENP and Borealin mutants (CPC_I1-190SB_, CPC_I1-190_6ASB_, CPC_I1-190SBΔloop_, CPC_I1-58SB_ and CPC_I1-190SB10-end_; Fig. S3C and S4F) with H3T3ph nucleosomes reconstituted with either Widom 601 or centromeric α-satellite DNA and performed Micrococcal nuclease (MNase) assays (Fig. 5E). Our MNase data shows that the DNA wrapped around the H3T3ph histone octamer was protected in the presence of CPC_I1-190SB_, while removing the Borealin loop region (CPC_I1-190SBΔloop_) and either mutating the INCENP ‘RRKKRR’ motif (CPC_I1-190_6ASB_) or deleting the entire basic IDR of INCENP (CPC_I1-58SB_), all showed a clear decrease in DNA protection. We also observed a comparable decrease in DNA protection when we perturb the Borealin interaction anchoring CPC to the nucleosome acidic patch (CPC_I1-190SB10-end_).

To assess if CPC can exert similar protection against MNase activity in the context of chromatin, we performed the above described MNase assay with a 12-mer H3T3ph nucleosomal array (Fig. 5F and S4G). In the absence of CPC, 2 minutes of MNase digestion resulted in mono- and di-nucleosomes (Fig 5F). Addition of CPC_I1-190SB_ resulted in a significant protection with ∼15% of the nucleosomes still existing as an intact 12-mer (Fig 5F). Consistent with our mononucleosome MNase data, all Borealin and INCENP mutants showed reduced MNase protection activity (Fig 5F).

### CPC-nucleosome interaction is essential for centromeric chromatin protection

Abolishing the centromere enrichment of CPC either by blocking the H3T3ph (Haspin-mediated) and H2AT120ph (Bub1-mediated) pathways or by removing the Borealin loop region critical for chromosome association of CPC resulted in severe mitotic defects, highlighting the essential requirement of centromere association of CPC (Liu *et al*, 2013b; Jeyaprakash *et al*, 2007; Abad *et al*, 2019; Liang *et al*, 2020; Broad *et al*, 2020; Hadders *et al*, 2020). Interestingly, our finding that INCENP ‘RRKKRR’ mutant does not completely abolish CPC-NCP binding, but weakens the nucleosome stabilizing activity of CPC *in vitro*, provides an opportunity to assess the consequence of perturbing nucleosome stabilizing role of CPC on the centromere enrichment of CPC and accurate cell division. We generated RPE1 cell lines expressing mNeonGreen-Borealin in which endogenous INCENP could be replaced either with INCENP wild type or INCENP 6A mutant and performed Fluorescence Recovery After Photobleaching (FRAP) analysis in prometaphase cells (Fig. 6A, Fig. S5A and S5B). Cells expressing mNeon-Green-Borealin were photobleached at centromeres and the signal recovery was determined from time lapse images. FRAP analysis showed that fluorescence recovery of the CPC containing the INCENP 6A mutant is faster compared to the CPC containing INCENP WT, indicating that perturbing the INCENP ‘RRKKRR’ contribution to CPC-nucleosome binding leads to less stable centromere association of CPC during prometaphase (Fig. 6A).

**Fig 6.**
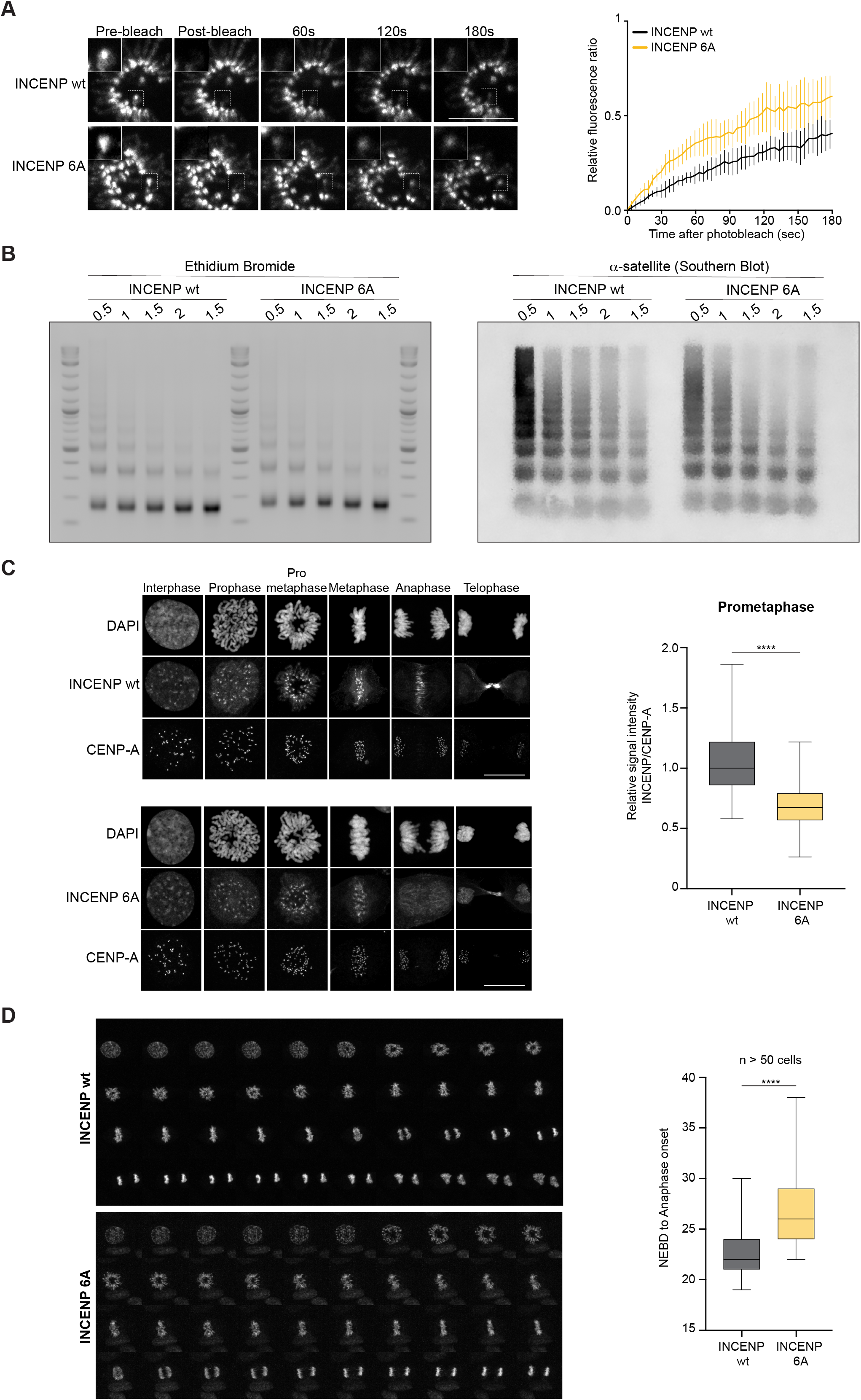
CPC-mediated protection of centromeric chromatin is crucial for accurate chromosome segregation. **(A)** (Left panel) FRAP analysis of WT and 6A mutant cell lines during prometaphase. The boxed area was bleached, and signal recovery is shown at the indicated time points. Scale bar, 10 μm. (Right panel) Quantification of FRAP experiments. Solid lines represent the means of quantitative FRAP measurements of Borealin-mNeonGreen in at least 10 prometaphase cells for each cell line (WT in black and 6A mutant in yellow). Light lines represent the standard error at each time point. **(B)** (Left panel) Representative agarose gel for MNase assay of chromatin, stained with ethidium bromide. (Right panel) Corresponding southern blot of the MNase assay with an alpha-satellite probe to blot for centromeric chromatin. The MNase concentrations from left to right are 0.125, 0.25, 0.5, 1, and 2 U/mL. **(C)** Representative fluorescence images for INCENP wt and INCENP 6A centromere localisation. DAPI was used for DNA staining. Scale bar, 10 μm. **(D)** Representative micrographs of live cell imaging experiments to assess the mitotic timing for INCENP wt and INCENP 6A mutant. Box-and-whisker plot showing the elapsed time (min) between nuclear envelope breakdown (NEBD) and anaphase onset/death for individual cells.

We then assessed whether the reduced stable centromere association of the CPC impacts centromeric chromatin stability by performing MNase digestion of chromatin on mitotic lysates of RPE1 cell lines expressing either INCENP wt or INCENP 6A. Consistent with our *in vitro* MNase data (Fig. 5F), in cells expressing the INCENP 6A mutant, the centromeric chromatin is more sensitive to MNase digestion during prometaphase (Fig. 6B and S5C).

In agreement with previous reports (Serena *et al*, 2020), INCENP 6A mutant failed to translocate from centromeres to the central spindle (Fig. 6C). Notably, we observed that the accumulation of the INCENP 6A mutant at centromeres during prometaphase was significantly reduced compared to the wild type (Fig. 6C). In addition, the live cell imaging experiments carried out with the RPE1 cell lines showed a significant delay in mitotic progression in cells expressing the INCENP 6A mutant (Fig. 6D).

Overall, this cellular data, along with the *in vitro* biochemical analysis, highlights the requirement of the chromatin stabilizing role of CPC in ensuring accurate cell division.

## Discussion

Efficient enrichment of CPC at the inner centromere during early stages of mitosis is essential for ensuring faithful chromosome segregation. Over the years numerous studies have investigated CPC localization and function (Klebig *et al*, 2009; Kelly *et al*, 2010b; Wang *et al*, 2010; Yamagishi *et al*, 2010; Kawashima *et al*, 2010; Tsukahara *et al*, 2010; Jeyaprakash *et al*, 2011; Trivedi & Stukenberg, 2016; Abad *et al*, 2019; Serena *et al*, 2020; Abad *et al*, 2022). However, how the different components of CPC collectively contribute towards chromatin interaction, and whether CPC has any Aurora B kinase independent role in influencing chromatin organization and/or integrity, are yet to be understood.

In this study, we aimed to answer these long-standing questions by characterizing CPC-nucleosome interaction employing an integrative structure-function approach combining structural, biochemical and cell-based approaches. The high-resolution cryo-EM structure of CPC bound to H3T3phNCP reported here identifies stable interactions of CPC with nucleosome acidic patch and the nucleosomal DNA entry-exit site, mediated via a highly basic Borealin N-terminal tail and the triple helical bundle formed by Borealin, Survivin and INCENP, respectively. Crosslinking/MS analysis and further biochemical characterization revealed dynamic nucleosome interactions involving the Borealin loop and INCENP basic IDR, underscoring their crucial role for efficient CPC-NCP binding. We previously showed that Borealin dimerization domain contributes to efficient nucleosome binding, but how it does so remained unclear (Abad *et al*, 2019). Our observation that two copies of CPC engage with both faces of the nucleosome (related by a two-fold symmetry) suggests the potential contribution of Borealin dimerization in facilitating this interaction.

Strikingly, the occupation of two CPCs on both faces of the nucleosome, while facilitating the interaction of CPC triple helical bundle with nucleosome DNA entry-exit sites, completely occludes the H2A C-terminal tail, making it inaccessible to its binding partners. Moreover, the H2A C-terminal tail residue T120 directly interacts with the CPC triple helical bundle (Fig S5D). Phosphorylation of H2A Thr 120 by Bub1 is known to be essential for Sgo1 localization to centromeres and for Sgo1’s role in protecting cohesion and preventing premature sister chromatid separation (Kitajima *et al*, 2005; Kawashima *et al*, 2010; Liu *et al*, 2013a; Hengeveld *et al*, 2017; Broad *et al*, 2020). Remarkably, the NCP binding mode of the CPC occludes the accessibility of the histone H2A tail for Sgo1 binding (Fig. S5D). These observations, together with our earlier study showing that the CPC binding of both Sgo1 and H3T3ph requires the same binding pocket on the Survivin BIR domain, suggest that the H3T3ph-mediated and the H2AT120ph-mediated CPC binding are mutually exclusive and likely to be restricted spatially (H3T3ph mediate inner centromere vs H2AT120 mediated kinetochore proximal) and possibly also temporally (Liang *et al*, 2020; Broad *et al*, 2020; Hadders *et al*, 2020; Abad *et al*, 2022; Cairo *et al*, 2023).

Haase et al., previously reported an Aurora B kinase independent non-catalytic role of CPC in facilitating kinetochore assembly. Their work showed that centromere association of the CPC localization module was sufficient to rescue the inner kinetochore assembly defects observed upon CPC depletion (Ditchfield *et al*, 2003; Wang *et al*, 2011; Wheelock *et al*, 2017; Haase *et al*, 2017). Our data reported here show that CPC-nucleosome binding stabilizes nucleosomal array *in vitro* and centromeric chromatin *in vivo* by employing a multipartite interaction involving both stable and dynamic interactions. We speculate that this non-catalytic activity of CPC likely contributes to stabilizing and maintaining the correct local architecture of the centromeric chromatin crucial for functional kinetochore assembly. It is also conceivable that CPC and the associated inner centromere interaction network (involving Sgo1, HP1 and cohesin) might also contribute to CPC-mediated tension sensing and error-correction.

Note: A recent study on biorxiv, Ruza et al, 2025, reports a cryo-EM structure of CPC bound H3T3ph nucleosome. While the mode of nucleosome binding by CPC broadly agrees with observations reported here, their structure shows just one copy of CPC bound to nucleosome, possibly because their CPC construct lacks more than two-thirds of the C-terminal region of Borealin (including the Borealin loop and dimerization domain).

## Materials and Methods

### Protein expression and purification of CPC constructs

The vectors used to express and purify CPC were: pMCNcs-INCENP_1-58_ for untagged INCENP_1-58_, pEC-S-CDF-His INCENP for His-tagged INCENP_1-190_, pRSET-GFP-Survivin for GFP-tagged full length Survivin and pETM-K-TEV-His-Borealin (pETM vector, gift from C. Romier, Institute of Genetics and Molecular and Cell Biology, Strasbourg, France) for His-tagged Borealin full length, Borealin_10-end_ and Borealin_Δloop_.

CPC constructs were purified as described in (Abad *et al*, 2019) and (Abad *et al*, 2022). Briefly, the three vectors for INCENP, Survivin and Borealin constructs were co-transformed in BL21(DE3) pLysS cells. The bacterial cultures were then grown at 37 °C until an OD of 0.6-0.8 was reached. Cultures were then induced overnight at 18 °C with 0.35 mM IPTG. The cultures were then harvested by spinning at 4,000 xg at 4 °C for 15 minutes and the pellets were resuspended in lysis buffer containing 20 mM Tris pH 8.0, 500 mM NaCl, 35 mM imidazole and 2 mM β-Mercaptoethanol, supplemented with complete EDTA-free cocktail tablets (Roche), 0.01 mg/ml DNase (Sigma-Aldrich), and 1 mM PMSF. The lysate was then sonicated at 40 % output for 3 rounds of 5 minutes (pulse: 2 on and 2 off), and clarified post sonication at 22,000 rpm for 50 mins. The supernatant was then loaded into a pre-equilibrated 5 ml His Trap Column (Cytiva) for affinity chromatography. The protein-bound column was washed with lysis buffer followed by chaperone buffer (20 mM Tris pH 8.0, 1000 mM NaCl, 35 mM Imidazole, 50 mM KCl, 10 mM MgCl_2_, 2 mM ATP and 2 mM β-Mercaptoethanol). The protein was then eluted using elution buffer (20 mM Tris pH 8.0, 500 mM NaCl, 500 mM Imidazole, 2 mM β-Mercaptoethanol) and dialyzed overnight in dialysis buffer (25 mM Hepes pH 8.0, 250 mM NaCl, 2 mM DTT) while cleaving with TEV and 3C proteases for affinity tag cleavage. The dialyzed samples were spun down and the supernatant was loaded onto a HiTrap SP HP (Cytiva) cation exchange column to separate the CPC_ISB_ stoichiometric complexes from other sub-complexes and tags. The fractions containing stoichiometric and pure CPC_ISB_ complex were pooled, concentrated, and run on a Superdex 200 Increase 10/300 (Cytiva) pre-equilibrated with 25 mM Hepes pH 8.0, 250 mM NaCl, 5 % Glycerol and 1 mM DTT. Protein fractions were pooled, concentrated and snap frozen for storage at -80oC.

### Protein expression and purification of Histones

Histones were purified based on the protocol from (Luger *et al*, 1999). For H2A, H2B and H3 histones, the vectors were transformed into BL21 pLysS, while H4 histone vector was transformed into BL21 Gold. The bacterial cultures were then grown at 37 °C until an OD of 0.5-0.7 was reached. They were then induced with 0.2 mM IPTG for 3 hours. The culture was harvested by spinning at 5,000 xg at RT for 10 minutes and the pellet was resuspended in ice-cold wash buffer containing 50 mM Tris pH 7.5, 100 mM NaCl, 1 mM EDTA, 1 mM Benzamidine and 1mM β-Mercaptoethanol. The resuspended cultures were snap frozen and thawed three times to lyse the cells. The lysates were then sonicated on ice with 60 % output for a total of 7.5 mins. The lysate was then clarified by spinning at 22,000 rpm for 60 mins at 4°C. After centrifugation, the pellet was homogenized using a dounce glass/glass homogenizer in wash buffer with and without 1% triton X-100. The pellet was then soaked in DMSO for 15 minutes and then homogenized once more in unfolding buffer containing 20 mM Tris pH 7.5, 7000 mM GuHCl, and 10 mM DTT. One more round of sonication was done for 15 s at 40 % output, and the lysate was stirred at RT for an hour, and then centrifuged at 23,000g for 20 min at 4°C. The supernatant was then dialyzed in Urea Buffer containing 10 mM Tris pH 8, 100 mM NaCl, 7000 mM Urea, 1 mM EDTA, and 5 mM β-Mercaptoethanol overnight in three rounds. After dialysis, the samples were centrifuged at 45,000 xg for 30 mins at 4°C. The samples were once again sonicated for 10 seconds at 40 % output, and filtered through a Millipore combined glass/PVDF filters (SLHVM25NS) to clear any remnant DNA contamination. The sample was then loaded onto a HiTrap Q column and HiTrap SP HP (Cytiva) cation exchange column sequentially to get pure histone fractions. Histones were eluted from the HiTrap SP column and the fractions containing histones were pooled, dialyzed against water supplemented with acetic acid and lyophilized for storage in -80 °C. The tail-less histone H3 used for native ligations (H3T32C) was purified using the same procedure with the following changes: pH 7 was used for all the buffers and the Urea dialysis buffer contained 10 mM Tris pH 8, 50 mM NaCl, 7000 mM Urea, 1 mM EDTA, and 5 mM β-Mercaptoethanol.

### Native Ligation of H3T3ph Histone tail

The native protocol protocol is based on (Bartke *et al*, 2010; Cistrone *et al*, 2019) and subsequently modified based on advice from Philipp Voigt (Babraham Institute). The modified histone tail was ordered from Peptide Synthetics Ltd. Initially the lyophilized H3 tail-less histone (H3T32C) was dissolved in methoxyamine buffer containing 6000 mM GuHCl, 400 mM methoxyamine, and 20 mM TCEP with a pH of around 3-4 to a concentration of around 32 mg/ml. The dissolved histone was then incubated at RT overnight, after which the methoxyamine was removed by dialyzing for 2 hours in ligation buffer (6M GuHCl, 200 mM Na2HPO4 pH 6.5-7, 20 mM TCEP). The modified histone tail peptide was dissolved in ligation buffer supplemented with 400 mM MPAA, and the pH adjusted to 4-5. The peptide solution was then mixed with the histone solution, and after adjusting the pH to around 7, the mixture was incubated at 23°C at RT for 2-3 days. Following the ligation, the mixture was dialyzed against Urea buffer (10mM Tris pH 8, 100 mM NaCl, 7000 mM Urea, 1 mM EDTA, and 1 mM DTT) overnight. Resource S column was used to separate the ligated histones from the unligated ones and the excess peptide. Fractions with ligated histones where then pooled, dialyzed overnight against water and lyophilized for storage.

### Nucleosome preparation

Lyophilized histones were resuspended in unfolding buffer, 7000 mM Guanidine HCl, 20 mM Tris pH 7.5, 10 mM DTT, and spun down at 14,000 xg for 5 minutes. After checking the concentration using Bradford, H3:H4:H2A:H2B were mixed in the ratio 1:1:1.1:1.1 and run on a SDS PAGE gel to confirm the stoichiometry. The histone mixture was then dialyzed for 2 hours in refolding buffer, 10 mM Tris pH 8, 2000 mM NaCl, 1 mM EDTA, 5 mM β-Mercaptoethanol, using an 8 kDa molecular weight cut-off dialysis membrane. Following 2 hours, the refolding buffer was replaced with fresh buffer, and the mixture was dialyzed overnight. A final dialysis was done for 2 hours with fresh refolding buffer, and the octamer was then spun down, filtered and run on a Superdex 200 Increase 10/300 (Cytiva) pre-equilibrated with refolding buffer. The stoichiometric fractions were pooled, snap frozen and stored at -80oC.

For Widom DNA, after PCR amplification of the widom 147 bp sequence from a pBS-601 Widom vector, Resource Q was carried out to purify the DNA. The Widom 601 147 bp sequence used was: ACAGGATGTATATATGTGACACGTGCCTGGAGACTAGGGAGTAATCCCCTTGGCGGTTAAAACGCGGGGGACAGCGCGTACGTGCGTTTAAGCGGTGCTAGAGCTGTCTACGACCAATTGAGCGGCCTCGGCACCGGGATTCTCCAG. For the α-satellite DNA, 10 midi preps were performed to purify the pUC57 containing 6 x 147 bp α-satellite DNA (kindly gifted by Prof. Ben Black; (Falk *et al*, 2015)), followed by an EcoRV digestion before loading the sample into a Resource Q column to obtain clean α-satellite DNA. The histone octamers were then titrated with the widom/α-satellite DNA by dialyzing the mixture in high salt refolding buffer (10 mM Tris pH 8.0, 2000 mM NaCl, 1mM EDTA and 5 mM β-Mercaptoethanol). The salt concentration was slowly brought down by adding low salt TE buffer (10 mM Tris pH 8.0, 50 mM NaCl and 1 mM EDTA) to facilitate the wrapping of the widom/α-satellite DNA around the octamer as described in (Luger *et al*, 1999). The titration was initially done on a small scale to determine the optimal ratio of Octamer:DNA for nucleosome reconstitution. After determining the correct ratio, large scale dialysis was done to obtain the nucleosomes used in our assays.

### Preparation of nucleosome arrays

The DNA template containing twelve tandem repeats of 167-bp 601 Widom sequence was kindly provided by Dr Dongyan Tan (originally from Dr. Craig Peterson). Large amounts of plasmid were prepared using a Macherey-Nagel PC 10000 kit (740593), following the manufacturer’s instructions. The desired 12-mer positioning sequence was cut out by an AvaI (NEB, R0152S) and HindIII (NEB, R3104S) restriction digest. The rest of the vector was fragmented using restriction enzymes DraI (NEB, R0129L). The 12-mer nucleosomal array DNA was isolated by size-exclusion chromatography using a Sephacryl-S500 16/60 column (GE Healthcare). Reconstitution into nucleosome arrays was performed by mixing array DNA at a final concentration of 0.3 μg/μl with 3 times molar excess of H3T3ph histone octamer (in proportion to available binding sites) and 0.1 mg/ml BSA (NEB, B9200S) in the high-salt buffer HEN2000 (10 mM HEPES, pH 8; 0.1 mM EDTA; 2 M NaCl). For array assembly tenfold volume of buffer HE (10 mM HEPES, pH 8; 0.1 mM EDTA) was added over 24 h at 4 °C. Final assembly took place by dialyzing against fresh low-salt buffer HN50 (10 mM HEPES, pH 8; 50 mM NaCl) for at least 3 h. Arrays were then precipitated by 5 mM Mg for 15’ at RT, spun down and the pellet was resuspended in 10 mM HEPES, pH 8; 5 mM KCl yielding a concentrated high-quality array.

### CPC-NCP complex formation using SEC

SEC experiments with recombinant purified proteins were performed using a MicroAKTA system (Cytiva), with Superose 6 5/150 (Cytiva) pre-equilibrated in 20 mM Hepes pH 8, 250 mM NaCl, 1 mM EDTA, and 1 mM DTT. 5 times molar excess of CPC_ISB_ was incubated with H3T3ph mono-nucleosomes and 15 mM ATP for 30 mins on ice before injection. UV 280 nm and 260 nm wavelengths were monitored to study complex formation, and 0.05 ml fractions were collected and run on SDS PAGE gels.

### CryoEM sample preparation and data collection

CPCI1-190 SB-H3T3ph mono-nucleosome complex was prepared by SEC as mentioned above. Fractions containing the stoichiometric complex were pooled and diluted to a final DNA concentration of 115 ng/ul right before grid preparation. 4.5 μl of sample was applied onto Quantifoil R2/2 300-mesh grids previously glow-discharged for 7 s in the presence of air using a current of 20 mA. Grids were blotted for 1.5 s at 10 °C and 95 % humidity using a Leica EM GP plunger (Leica). 22,065 micrographs were collected using a Titan Krios transmission electron microscope operating at 300 keV equipped with a Falcon4i direct electron detector and a Selectris X Energy Filter (energy slit width of 5 eV) (ThermoFisher). Smart EPU software (ThermoFisher) was used for the automated Data collection, MotionCor2 was used to perform motion correction of the movies (Zheng et al, 2017). The parameters used for data collection are provided in Table 1.

Motion corrected micrographs were processed using CryoSPARC v4.7.0 (Punjani *et al*, 2017). AI-based picking tool Topaz (Bepler *et al*, 2019a, 2019b) was initially used for particle picking with a previously trained nucleosome model, and the particles were extracted with a box size of 384 px, after which multiple rounds of 2D classification were performed to filter out junk particles and select the 2D classes presenting high resolution features. 1.6 million particles were used to generate several ab initio models and subsequent heterogeneous refinement, from which maps showcasing nucleosomes with well-defined bound CPC were selected. Further 3D classification followed by non-uniform refinement (Punjani *et al*, 2020) yielded three maps displaying three different CPC orientations at a resolution of 2.83 Å, 2.85 Å, and 2.86 Å (Fourier shell correlation (FSC) = 0.143). 3D variability was performed combining the particles corresponding to the three final maps to study the dynamicity of CPC binding to nucleosomes (Movie1). For each of the three final 3D volumes, AI-based DeepEM Enhancer sharpening was performed (Sanchez-Garcia *et al*, 2021). A crystal structure of the CPC triple helical bundle (PDB: 2QFA) and of human NCP (PDB: 2CV5) were initially fitted into the cryo-EM map using the rigid body fitting built-in tool in ChimeraX (version 1.9). Model building was done by performing iterative rounds of real-space refinement in Phenix and Coot. Interactions were analysed using PDBePISA (https://www.ebi.ac.uk/pdbe/pisa/).

### AFM sample preparation and imaging

Mono-nucleosomes for AFM was prepared by wrapping H3T3ph histone octamers with a 486 bp DNA construct comprising the W601 nucleosome positioning sequence flanked by a short (106 bp) and long (233 bp) extra-nucleosomal DNA arm: caatcaaactggctcgtcgcgaattggagctccaccgcggtggcggccgctcgatctagtactagtggcatgtcagctgcaggaattcgagctcaacgtgcaatccctggagaatcccggtgccgaggccgctcaattggtcgtagacagctctagcaccgcttaaacgcacgtacgcgctgtcccccgcgttttaaccgccaaggggattactccctagtctccaggcacgtgtcagatatatacatcctgtcgacgacacgggtgatcgactagttctagagcgatctagtatcgatcactcttttgttccctttagtgagggttaatttcgagcttgcgatctagtcgatactacgcgtaatcatggtcatagctgtttcctgtgtgaaatgatctacttgttatccgctcacaattccacacaacatacgagccggaaaagaaataaagatctagactactgcataaagtgtaaagcctggg (Genart strings DNA fragment, Geneart, Thermo Fisher Scientific). The final nucleosome sample contained approximately 50% free DNA, as was desirable for the experiment.

Poly-L-lysine-coated mica was prepared by drop-casting 20 μL poly-L-lysine (Merck; 0.01% w/v in autoclaved milliQ water) on freshly cleaved muscovite mica (SPI Supplies) for 30 s and subsequently rinsing the surface with 20 mL of milliQ water before drying with a gentle stream of filtered N2 gas (Vanderlinden *et al*, 2014). Mononucleosome samples (at 0.75 ng/uL of DNA) were incubated in aqueous buffer comprising 200 mM NaCl, 20 mM HEPES-KOH (pH 7.4), and 1mM DTT for 10 min on ice. The sample was then deposited on the poly-L-lysine-coated muscovite mica for 15 s and rinsed with 20 mL milliQ water before drying with a gentle stream of filtered N_2_ gas.

A Nanowizard 4 XP AFM (JPK, Berlin, Germany) was used in tapping mode with silicon tips (FASTSCAN-A; drive frequency, 1400 kHz; Bruker) over fields of view of 6 × 6 μm at 4,096 × 4,096 pixels and captured at line rates of 3 Hz. For each condition two independent data sets were recorded. The nucleosome samples for each data set were prepared in independent nucleosome reconstitutions.

### AFM image analysis

To process the raw topographic data, we used SPIP software for plane correction that included plane-fitting with a 3rd degree polynomial, and line-by-line correction with a 4th degree polynomial. The processed AFM topographs were further analyzed through a previously published, open-source automated image analysis pipeline (Konrad *et al*, 2021b); available at GitHub: https://github.com/SKonrad-Science/AFM_nucleoprotein_readout). In brief, image analysis starts with molecule detection and classification. To this end, a Gaussian filter and a background subtraction are applied followed by skeletonization whereby molecules are trimmed to a one-pixel-wide backbone. This skeleton is used for classification: bare DNA has exactly two endpoints in its skeleton and no branchpoints, while mono-nucleosomes have exactly two endpoints and two branchpoints. In a second analysis step, the tip shape and size is estimated from the skeleton of a subset of bare DNA molecules. Using the estimated tip shape, image erosion is applied to minimize tip dilation effects. Last, the classified molecules are analyzed with respect to the structure parameters arm lengths, volume, and opening angle for nucleosomes and contour length for bare DNA. The structure parameters are exported as text files and used for plotting and statistical analysis in Python. To quantify the fraction of fully wrapped nucleosomes in an ensemble, we use the 2 D heatmap of wrapped length versus opening angles (Figure 4c, Supporting Figure S3d). In this plot, we identify the two dominant local maxima (corresponding to the most probable fully wrapped respectively partially unwrapped conformations). Along the line connecting these maxima the valley point was found. Then, the minimum was detected inside the rectangle defined by the local maxima as opposing vertices. This minimum and valley point define the separation line (green dotted line in Figure 4c): nucleosomes are said to be fully wrapped if they fall above it.

### Micrococcal Nuclease assay

For mono-nucleosomes with Widom 601 and α-satellite DNA, 1 μg of nucleosome with or without CPC_ISB_ were incubated on ice for 1 hour. 0.25 μl of MNase (NEB M0247S) was then added to the mix and made up to 50 μl with the reaction buffer (50 mM Tris-HCl pH 8, 100 mM NaCl, 3 mM CaCl2, 1 mM DTT), and incubated at RT for 10 mins. The reaction was then quenched using 10 μl of 500 mM EDTA and 20 μg of proteinase K was added per reaction and incubated at 50 °C for 30 mins to digest the proteins. The DNA was then purified using Minelute Qiagen kit, and run on an Agilent 2100 Bioanalyzer using a High Sensitivity DNA assay kit.

For nucleosomal arrays, 500 ng of array with or without CPC was digested with 0.025 U of MNase (Sigma-Aldrich, 5386-50U) in buffer containing 5 mM CaCl_2_. The reaction was stopped using STOP-buffer (50 mM Tris pH 7.5, 4% SDS, 0.5 mM EDTA, pH 8.0) at different incubation time points. Afterward samples were incubated for 1 h at 37 °C with Proteinase K (Thermo Fisher Scientific, EO0491) and digested DNA was isolated using the NEB Monarch PCR & DNA Cleanup Kit (NEB, T1130L). The isolated DNA fragments were separated using 1.5% (Biozym, 840004) agarose gel electrophoresis with 1×TAE as running buffer (120 V, 1.5 h) and stained with SybrSafe. Gels were imaged using a GE Healthcare Typhoon FLA9000 imager.

### EMSA

20 nM mono-nucleosomes were incubated for an hour with increasing amounts of CPC constructs on ice in EMSA reaction buffer containing 10 mM Tris pH 7.5, 100 mM/250 mM NaCl, 1 mM MgCl2, and 1 % Glycerol supplemented with 0.1 mg/ml BSA. The complex was then run on a 5.2 % polyacrylamide Tris-Glycine gel in 1X Tris-Glycine buffer at 100V for 1h 40min at 4 °C. The gel image was analyzed using ImageJ. Statistical significance of the difference between the % binding was established by a Kruskal–Wallis test with Dunn’s multiple comparisons test using Prism 7.0.

### Crosslinking/MS

Crosslinking experiments for the CPC_I1-190SB_-H3T3ph NCP complex were performed using EDC with N-hydroxysulfosuccinimide (Thermo Fisher Scientific). 3 μg of NCP was crosslinked with 2.5 molar excess of CPC_I1-190SB_ using either 30 or 60 μg of EDC and either 66 or 132 μg of N-hydroxysulfosuccinimide in 20 mM Hepes, pH 8, 200 mM NaCl, 1 mM EDTA, and 2 mM DTT for 2 hours at room temperature. The cross-linking reaction was quenched by the addition of 100 mM Tris-HCl, and crosslinking products were briefly resolved using 4–12% Bis-Tris NuPAGE (Thermo Fisher Scientific). Bands were visualized by short Instant Blue staining (Thermo Fisher), excised, reduced with 10 mM DTT for 30 min at room temperature, alkylated with 5 mM iodoacetamide for 20 min at room temperature, and digested overnight at 37 °C using 13 ng/μl trypsin (Promega). Digested peptides were loaded onto C18-Stage-tips (Rappsilber *et al*, 2007).

Liquid chromatography with tandem mass spectrometry (LC-MS/MS) analysis was performed as previously described (Abad *et al*, 2022) using an Orbitrap Fusion Lumos Tribrid Mass Spectrometer (Thermo Fisher Scientific) applying a “high-high” acquisition strategy. Peptide mixtures were injected for each mass spectrometric acquisition and separated on a 75 µm × 50 cm PepMap EASY-Spray column (Thermo Fisher Scientific) fitted into an EASY-Spray source (Thermo Fisher Scientific) and run at a column temperature of 50 °C. Mobile phase A consisted of water and 0.1 % vol/vol formic acid, while mobile phase B consisted of 80 % vol/vol acetonitrile and 0.1 % vol/vol formic acid. Peptides were loaded at a flow-rate of 0.3 μl/min and eluted at 0.2 μl/min using a linear gradient going from 2% mobile phase B to 40% mobile phase B over 159 (or 129) min, followed by a linear increase from 40 to 95% mobile phase B in 11 min. The eluted peptides were then loaded into the mass spectrometer, and MS data was acquired in the data-dependent mode with the top-speed option. For each 3 sec acquisition cycle, the mass spectrum was recorded in the Orbitrap with a resolution of 120,000. The ions with a precursor charge state between 3+ and 8+ were isolated and fragmented using higher-energy collisional dissociation (HCD) or electron-transfer/HCD (EThcD). The fragmentation spectra were recorded in the Orbitrap. Dynamic exclusion was enabled with single repeat count and 60 sec exclusion duration.

The mass spectrometric raw files were processed into peak lists using ProteoWizard (v3.0.20338; Kessner et al, 2008), and cross-linked peptides were matched to spectra using Xi software (v1.7.6.3; Mendes et al., 2018; https://github.com/Rappsilber-Laboratory/XiSearch) with in-search assignment of monoisotopic peaks (Lenz *et al*, 2018). The search parameters used were MS accuracy, 3 ppm; MS/MS accuracy, 10 ppm; enzyme, trypsin; cross-linker, EDC; max missed cleavages, 4; missing mono-isotopic peaks, 2; fixed modification, carbamidomethylation on cysteine; variable modifications, oxidation on methionine; and fragments b- and y-type ions (HCD) or b-, c-, y-, and z-type ions (EThcD) with loss of H_2_O, NH_3_, and CH_3_SOH. 1% on link level false discovery rate was estimated based on the number of decoy identification using XiFDR (Fischer & Rappsilber, 2017).

### UV Crosslinking/MS

3 μg of H3T3ph NCPs reconstituted with IR700-labelled 601 Widom 147bp DNA were incubated with 8 times molar excess of CPC_I1-190 SB_ for 1 h at 4 °C and UV-crosslinked using a Stratalinker UV 1800 crosslinker (Stratagene) with a 254 nM UV light bulb and a dose of 1 J/cm^2^. Samples were then loaded onto a 4-12 % Bis-Tris NuPAGE gel and subjected to MS analysis.

LC-MS/MS analyses were performed on Orbitrap Fusion™ Lumos™ Tribrid™ Mass Spectrometer (Thermo Fisher Scientific, UK) on a Data Independent Acquisition (DIA) mode, coupled on-line, to an Ultimate 3000 HPLC (Dionex, Thermo Fisher Scientific, UK). Peptides were separated on a 50 cm (2 µm particle size) EASY-Spray column (Thermo Scientific, UK), which was assembled on an EASY-Spray source (Thermo Scientific, UK) and operated constantly at 55oC. Mobile phase A consisted of 0.1% formic acid in LC-MS grade water and mobile phase B consisted of 80% acetonitrile and 0.1% formic acid. Peptides were loaded onto the column at a flow rate of 0.3 μL min-1 and eluted at a flow rate of 0.25 μL min-1 according to the following gradient: 2 to 40% mobile phase B in 150 min and then to 95% in 11 min. Mobile phase B was retained at 95% for 5 min and returned back to 2% a minute after until the end of the run (190 min).

MS1 scans were recorded at 120,000 resolution (scan range 350-1650 m/z) with an ion target of 5.0e6, and injection time of 20ms. MS2 was performed in the orbitrap at 30,000 resolution with a scan range of 200-2000 m/z, maximum injection time of 55ms and AGC target of 3.0E6 ions. We used HCD fragmentation with stepped collision energy of 25.5, 27 and 30. We used variable isolation windows throughout the scan range ranging from 10.5 to 50.5 m/z. Narrow isolation windows (10.5-18.5 m/z) were applied in the range 400-800 m/z and then gradually increased to 50.5 m/z until the end of the scan range. The default charge state was set to 3. Data for both survey and MS/MS scans were acquired in profile mode.

The DIA-NN software platform (Demichev *et al*, 2020) version 1.9.2. was used to process the DIA raw files and search was conducted against our in-house CPC protein complex. Precursor ion generation was based on the chosen protein database (automatically generated spectral library) with deep-learning based spectra, retention time and IMs prediction. Digestion mode was set to specific with trypsin allowing maximum of two missed cleavages. Carbamidomethylation of cysteine was set as fixed modification. Oxidation of methionine, and acetylation of the N-terminus were set as variable modifications. The parameters for peptide length range, precursor charge range, precursor m/z range and fragment ion m/z range as well as other software parameters were used with their default values. The precursor FDR was set to 1 %.

### Cell culture and cell lines

hTRET-RPE1 cells were cultured in DMEM supplemented with 10% FBS and 0.2 mM L-glutamine at 37℃ in a 5% CO2 environment. To generate an RPE1 cell line stably expressing OsTIR1(F74G), RPE1 cells were first subjected to CRISPR-Cas9-mediated knockout of the originally integrated puromycin resistance gene and then transfected with pMK444, which harbors the OsTIR1(F74G) gene (Addgene #121192), and pCMV-hyBase (PiggyBac transposase) using the Neon Transfection system (Thermo Fisher Scientific) as described (Hatoyama et al., 2024, EMBO). Transfected cells were selected with 1 µg/mL puromycin, and isolated as single clones. INCENP-mAID RPE1 cell line was generated from the OsTIR1 (F74G)-expressing RPE1 cells using CRISPR-Cas9-mediated homologous recombination. Briefly, approximately 500 bp homology arms corresponding to the INCENP gene were cloned into the donor vector pMK391 (Addgene #121192). A short guide RNA (sgRNA) targeting the C-terminus of INCENP was designed using the web-based prediction tool CHOPCHOP (https://chopchop.cbu.uib.no/) and replaced the originally inserted sgRNA in the sgRNA vector (Addgene #72833). These plasmids were transfected into OsTIR1(F74G)-expressing RPE1 cells using the Neon system. Transfected cells were selected with 50 µg/mL blasticidin, isolated as single clones, and validated as biallelic knock-ins using PCR and Western blotting. To generate doxycycline-inducible RPE1 cells expressing wild-type or 6A mutant INCENP, 3×Flag-tagged INCENP genes were inserted in the hROSA26 targeting vector harboring the TetOn 3G system (Addgene #114699) and transfected into the INCENP-mAID RPE1 cell line, along with the sgRNA plasmid targeting the hROSA26 locus (Addgene #105927). For the induction of gene expression from the hROSA26 locus and the degradation of the endogenous mAID-fused protein, cells were treated with 1 µg/mL doxycycline or 1 µM 5Ph-IAA for the indicated time. For mNeonGreen-tagging of Borealin, cells were transfected with a modified pMK286 donor plasmid (in which the mAID tag was replaced with mNeonGreen) and an sgRNA plasmid targeting the C-terminus of the Borealin gene, using the same procedure as described above.

### Immunofluorescence microscopy

Cells grown on coverslips were fixed with 2% paraformaldehyde in PBS for 10 minutes and permeabilized with 0.2% Triton X-100 in PBS for 10 minutes. After blocking with 3% BSA in PBS for 10 minutes, cells were incubated with primary antibodies diluted in 3% BSA in PBS. The following primary antibodies were used: anti-INCENP (1:3000; 29419-1-AP, Proteintech) and anti-CENP-A (1:500; 3-19, MBL). After incubation with secondary antibodies and counterstaining of DNA with 0.1 μg/mL DAPI, cells were mounted using ProLong Gold Antifade Mountant (Invitrogen). Images were acquired using a Plan-Apochromat 100×/1.46 NA oil immersion objective lens (Zeiss) on an LSM880 confocal laser scanning microscope (Zeiss).

### Antibodies used for Western Blotting

The following primary antibodies were used for Western blotting: anti-FLAG (1:5000; M2, Sigma) anti-INCENP (1:5000; 29419-1-AP, Proteintech) anti-Aurora B (1:500; 611083, BD Biosciences) anti-α-Tubulin (1:5000; clone B-5-1-2, #T6074, Millipore)

### FRAP

RPE1 cell lines were seeded onto chambered cover glasses (Thermo) the day before the experiment and, prior to imaging, were cultured at 37°C for at least 1 h in Leibovitz’s L-15 medium (Gibco) without phenol red, supplemented with 20 % fetal bovine serum (FBS; Sigma) and 20 mM HEPES (pH 7.45) (Hatoyama *et al*, 2024). For FRAP analysis, cells were cultured at 27 ℃ to minimize chromosome movement. The mobility of Borealin-mNeonGreen in each cell line was analyzed using a confocal microscope (LSM880, Carl Zeiss) equipped with a 63× objective lens (1.4 NA Plan Apochromat Oil DIC M27) for region-of-interest bleaching. Two images were acquired prior to bleaching (1% transmission of a 488-nm laser, 204.8 ms/frame with a 3 s interval, 256 × 256 pixels, 2 Airy unit pinhole, 10× zoom, 4-μm z-stack with 1-μm intervals). Borealin-mNeonGreen signals were bleached using 10 % and 100 % transmission of 405- and 488-nm lasers, respectively, followed by the acquisition of an additional 60 images using the same settings. The fluorescence intensity in the bleached region was quantified using Fiji software after z-stack projection to correct for centromere movement along the z-axis and background subtraction. Photobleaching during imaging was monitored and corrected before plotting the recovery curve. The recovery curve was plotted as relative values, setting the fluorescence intensities before and after bleaching as 1 and 0, respectively.

### Live-cell imaging

To analyze mitotic progression, live-cell imaging was performed using a confocal quantitative image cytometer CQ1 (Yokogawa). Cells were cultured overnight in a 24-well glass-bottom plate (Iwaki). DNA was stained with 100 nM SiR-DNA (Cytoskeleton, Inc.) for 1 h before image acquisition. Images were acquired every 1 min using a 60× objective lens. A series of projected images from five Z-sections at 1.5 µm intervals were analyzed. For data analysis, images were processed using Fiji software (http://fiji.sc).

### Southern Blot

To obtain mitotic cells, cells were cultured with 200 nM Palbociclib for 24 h, released for 12 h, and then treated with 0.1 μg/mL nocodazole for 4 h before being collected by shake-off. After washing with PBS, 5 × 10⁵ cells were suspended in a modified CSK buffer (0.1% Triton X-100, 100 mM NaCl, 10 mM PIPES [pH 7.0], 300 mM sucrose, 1 mM MgCl₂, 5 mM CaCl₂, 1 mM EGTA, and a protease inhibitor cocktail [Roche]) containing the indicated concentration of micrococcal nuclease (SIGMA) and incubated for 5 min at 37°C. The reaction was stopped with 50 mM EDTA, followed by treatment with RNase A and a protease inhibitor. The digested DNA was then purified by phenol/chloroform extraction and ethanol precipitation. The DNA was electrophoresed on a 1.2% agarose gel in TAE buffer. After electrophoresis, the gel was denatured in 0.5 M NaOH with 1.5 M NaCl, neutralized in 0.5 M Tris (pH 7.5) with 1.5 M NaCl, and equilibrated in 20× SSC. The DNA was then transferred onto a Hybond-N+ membrane (Amersham) by capillary blotting overnight. Hybridization and probe preparation were performed using the ECL Direct Nucleic Acid Labeling and Detection System (Cytiva). Alpha-satellite DNA fragments were amplified using the following primers (Chan *et al*, 2012) and monomeric unit was purified.

Forward: 5′-CTCACAGAGTTGAACCTTCC-3′

Reverse: 5′-GAAGTTTCTGAGAATGCTTCTG-3′

(Ref: Active transcription and essential role of RNA polymerase II at the centromere during mitosis)

## Acknowledgements

T.H. lab is supported by the Japan Society for the Promotion of Science (JSPS) Grant-in-Aid for Scientific Research (24H01381, 24H02286 [to R.-S. N.]; 24H02283, 22H04996 [to T.H.]), and by JST, CREST Grant Number PMJCR21E6, Japan [to T.H.]. This work was supported by funding for the Wellcome Discovery Research Platform for Hidden Cell Biology [226791] and we gratefully acknowledge support from the Proteomics core and the Atomic Force Microscopy facility. M.D.W.’s work is supported by the Wellcome Trust (210493), Medical Research Council (T029471/1). K.P.H. and his team are funded by Deutsche Forschungsgemeinschaft TTR237 and SFB1361. A.A.J. and his team are supported by the Wellcome Trust (senior research fellowship 202811), the European Union (ERC advanced grant CHROMSEG 101054950), and the Medical Research Council (MRC grant MR/X001245/1). The Wellcome Centre for Cell Biology is supported by core funding from the Wellcome Trust (grant 203149). A.G. is supported by the Darwin Trust of Edinburgh. Views and opinions expressed are, however, those of the authors only and do not necessarily reflect those of the European Union or the European Research Council. Neither the European Union nor the granting authority can be held responsible for them.

## Data availability

Cryo-EM structure and model are deposited in Protein Data Bank (PDB: https://www.rcsb.org/) under the following accession codes: PDB ID 9R77, Extended PDB ID pdb_00009R77, EMD-53735.

## Author Contributions

Conceptualization: A.A.J.; Funding acquisition: K.P.H., M.D.W., W.D.V., T.H., A.A.J.; Investigation: A.G., M.A.A., R-S.N., P.S-P., L.C.D., M.L., C.S., C.C.P., A.A.J.; Methodology: A.G., M.A.A., R-S.N., A.A.J.; Software: J.R., W.D.V.; Writing – original draft: A.G., M.A.A., A.A.J.; Writing – review & editing: A.G., M.A.A., R-S.N., P.S-P., L.C.D., M.L., C.S., W.D.V., A.A.J.

