## Supplementary Information for "Chromatin Protection by the Chromosomal Passenger Complex"

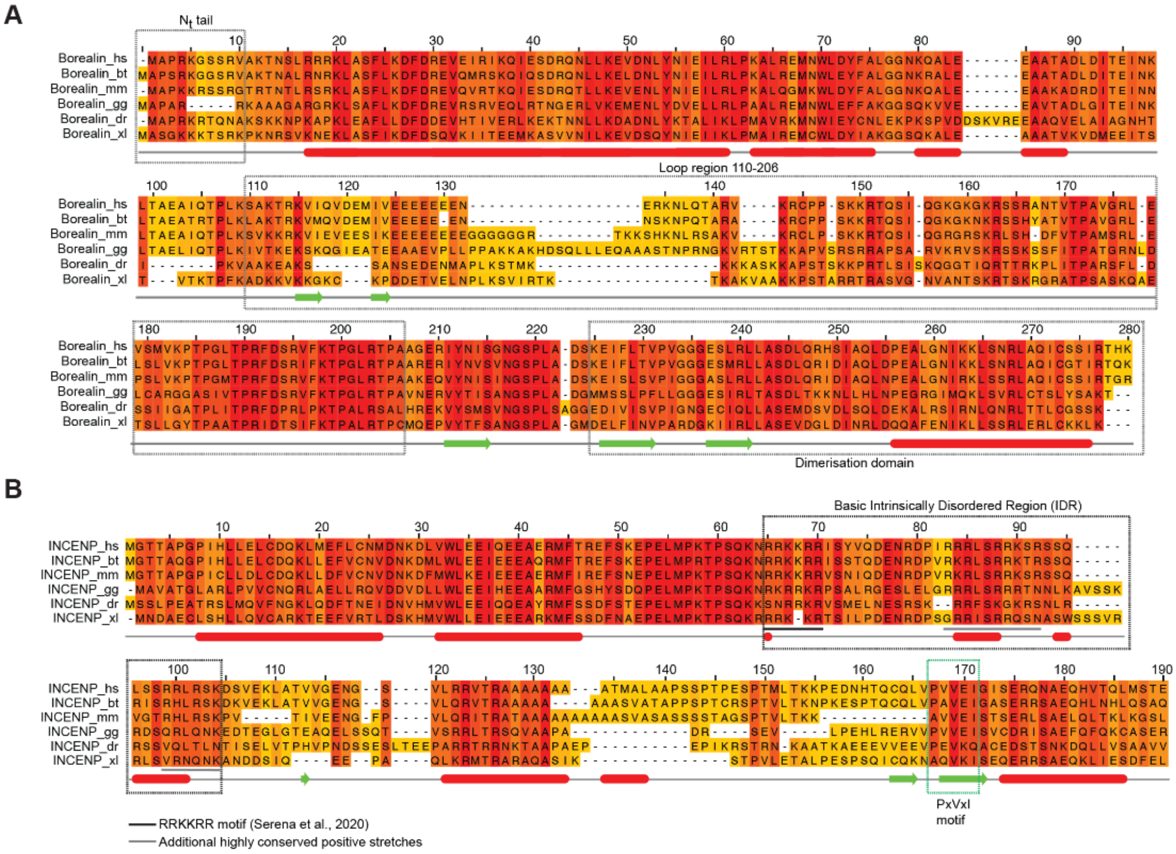
A
Nt tail
Borealin_hs
Borealin_bt
Borealin_mm
Borealin_gg
Borealin_dr
Borealin_xl
Loop region 110-206
Dimerisation domain
B
Basic Intrinsically Disordered Region (IDR)
INCENP_hs
INCENP_bt
INCENP_mm
INCENP_gg
INCENP_dr
INCENP_xl
PxVxI motif
RRKKRR motif (Serena et al., 2020)
Additional highly conserved positive stretches

**Figure S1**

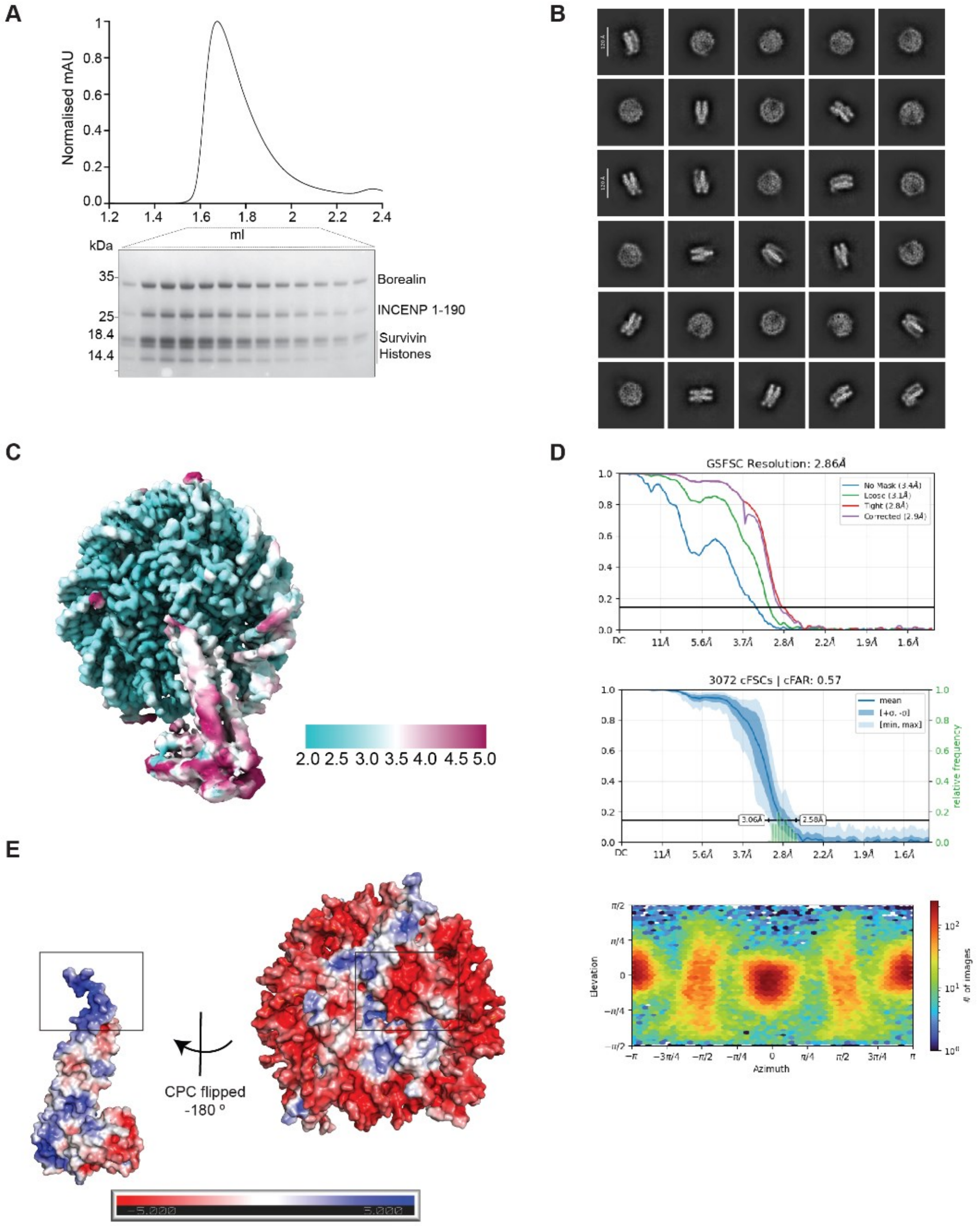
A
Normalised mAU
ml
kDa
35
25
18.4
14.4
Borealin
INCENP 1-190
Survivin
Histones
B
C
2.0 2.5 3.0 3.5 4.0 4.5 5.0
D
GSFSC Resolution: 2.86Å
No Mask (3.4Å)
Loose (3.1Å)
Tight (2.8Å)
Corrected (2.9Å)
3072 cFSCs | cFAR: 0.57
mean
relative frequency
3.06Å
2.58Å
Elevation
Azimuth
### of images
E
CPC flipped
-180 °
-5.000
5.000

**Figure S2**

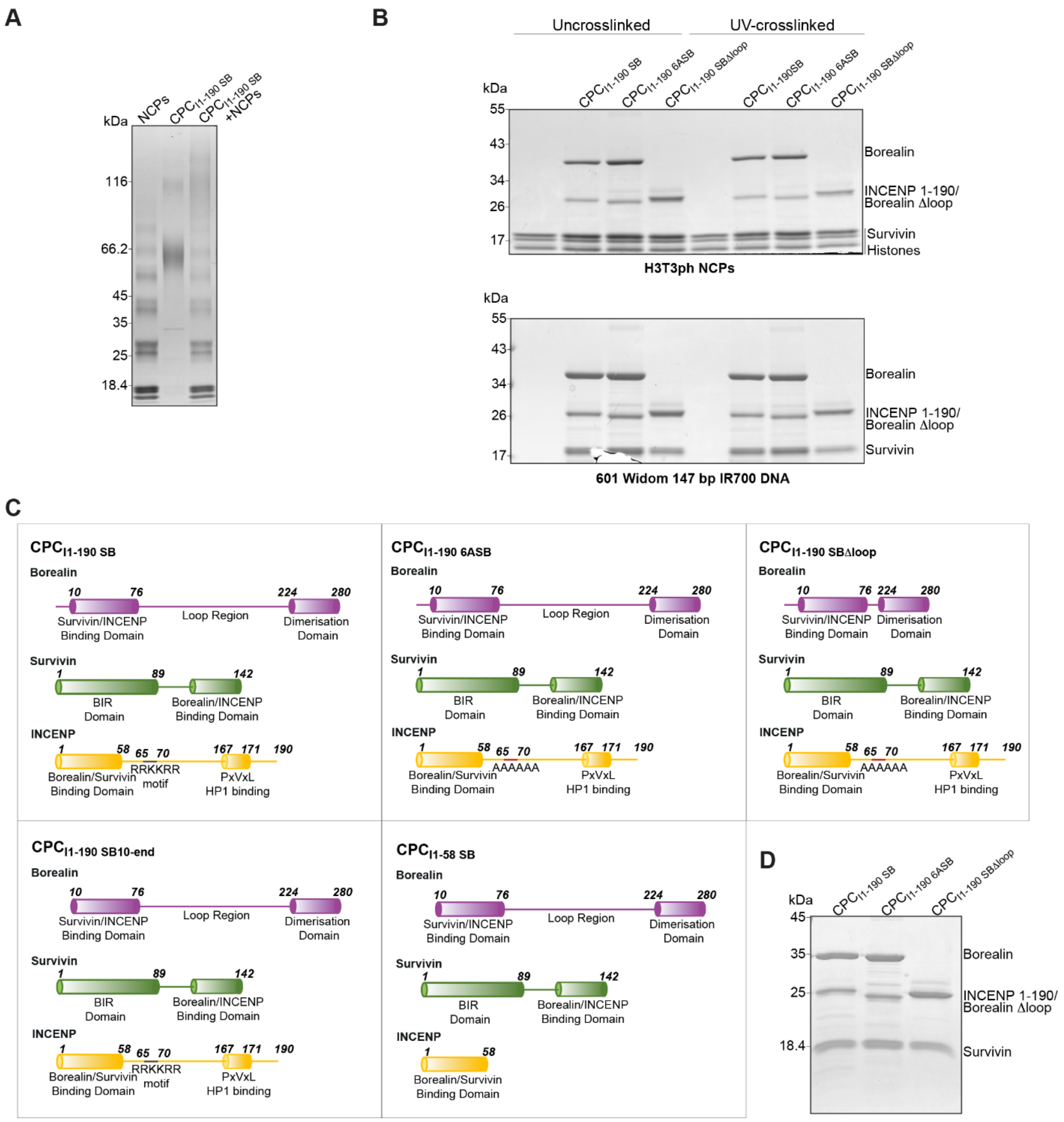
A
kDa
NCPs
CPC I1-190 SB
CPC I1-190 SB +NCPs
116
66.2
45
35
25
18.4
B
Uncrosslinked
UV-crosslinked
CPC I1-190 SB
CPC I1-190 6ASB
CPC I1-190 SBΔloop
Borealin
INCENP 1-190/
Borealin Δloop
Survivin
Histones
H3T3ph NCPs
601 Widom 147 bp IR700 DNA
C
CPC I1-190 SB
CPC I1-190 6ASB
CPC I1-190 SBΔloop
CPC I1-190 SB10-end
CPC I1-58 SB
Borealin
Survivin/INCENP Binding Domain
Loop Region
Dimerisation Domain
Survivin
BIR Domain
Borealin/INCENP Binding Domain
INCENP
Borealin/Survivin Binding Domain
RRKKRR motif
AAAAAA
PxVxL HP1 binding
D
Borealin
INCENP 1-190/
Borealin Δloop
Survivin

### Figure S3

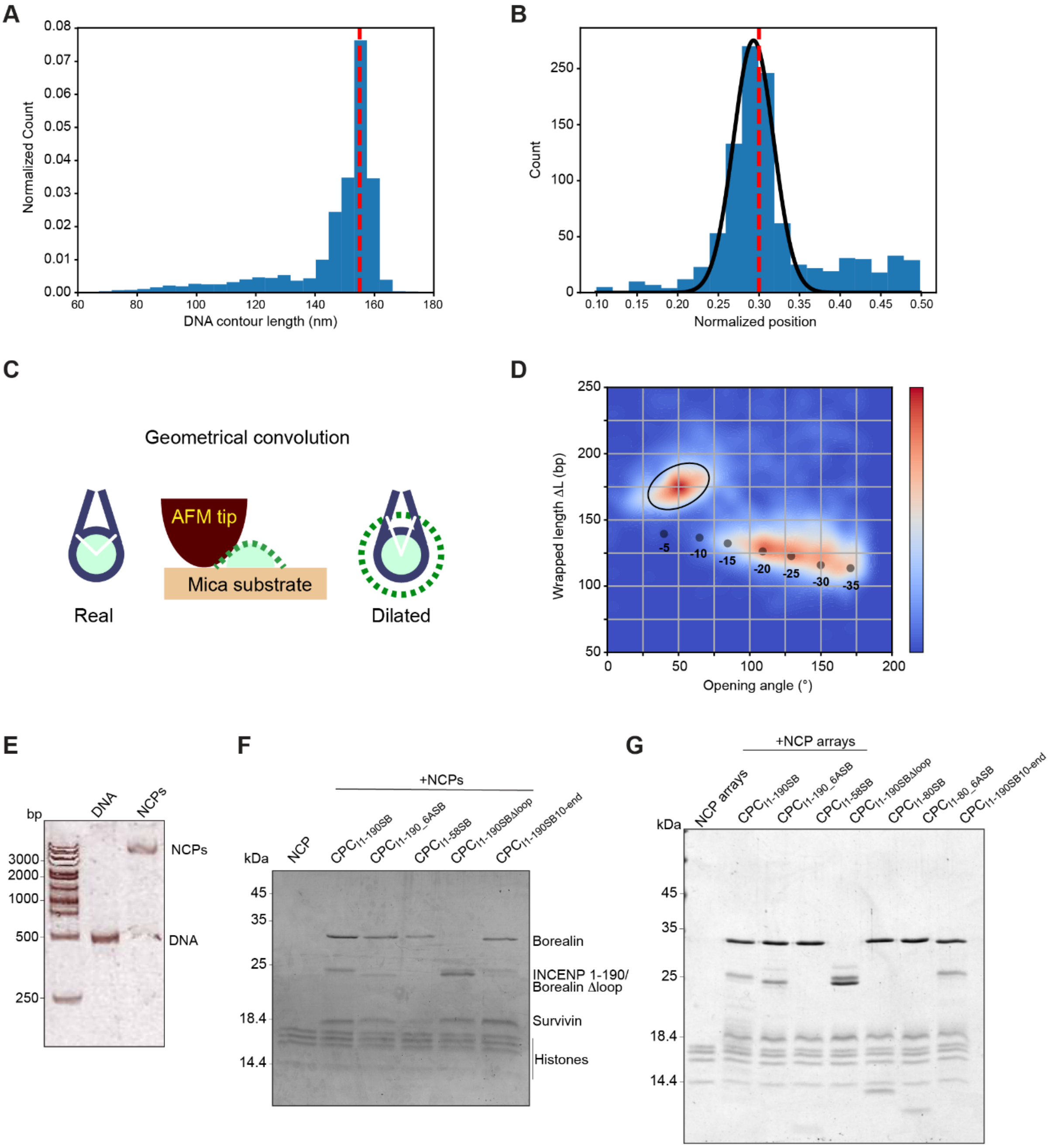
A
Normalized Count
DNA contour length (nm)
B
Count
Normalized position
C
Geometrical convolution
AFM tip
Mica substrate
Real
Dilated
D
Wrapped length ΔL (bp)
Opening angle (°)
-5
-10
-15
-20
-25
-30
-35
E
bp
DNA
NCPs
3000
2000
1000
500
250
F
+NCPs
kDa
NCP
Borealin
INCENP 1-190/
Borealin Δloop
Survivin
Histones
G
+NCP arrays
kDa
NCP arrays

**Figure S4**

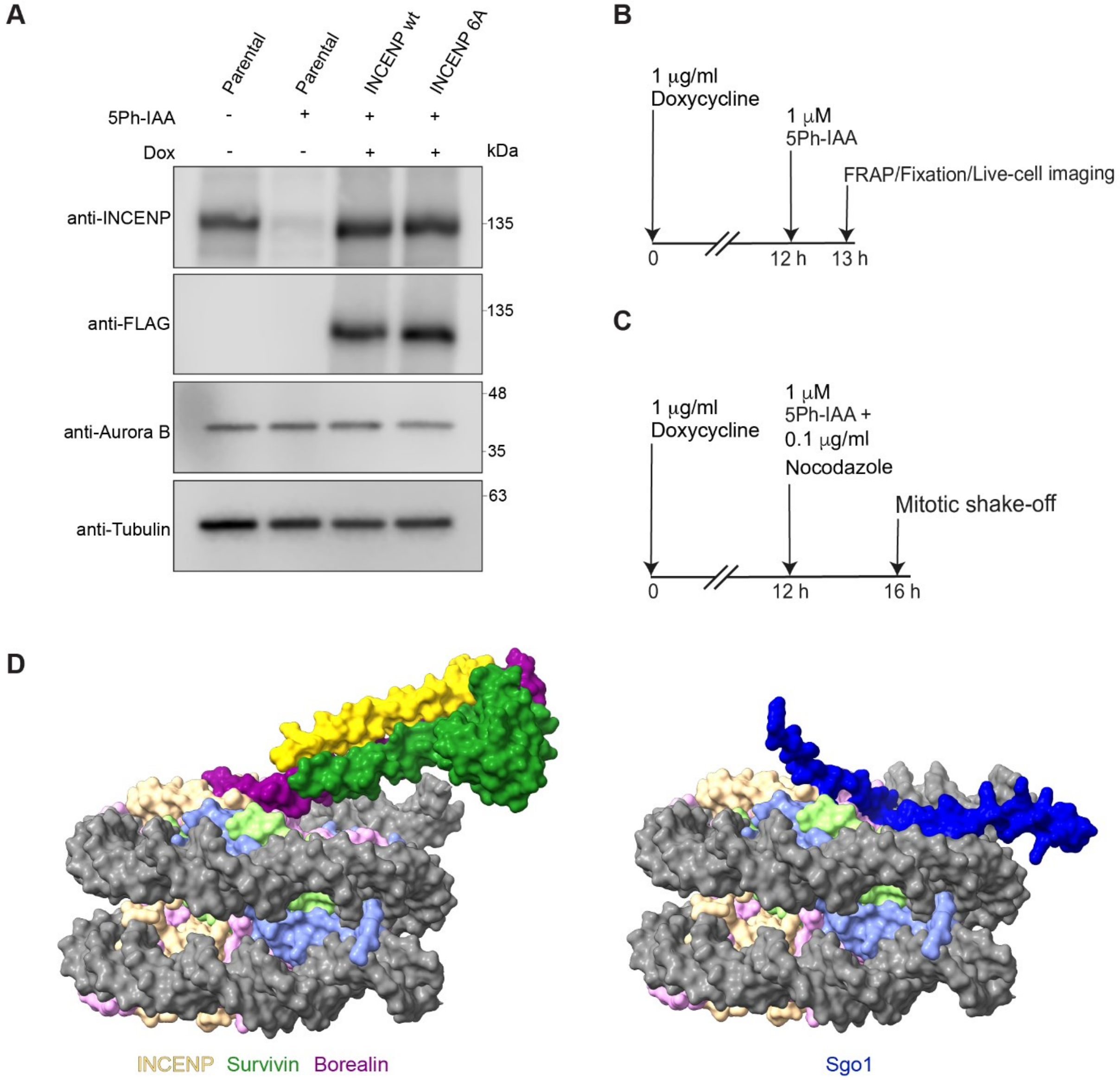
A
Parental
Parental
INCENP wt
INCENP 6A
5Ph-IAA - + + +
Dox - - + +
kDa
anti-INCENP
135
anti-FLAG
135
anti-Aurora B
48
35
anti-Tubulin
63
B
1 µg/ml Doxycycline
1 µM 5Ph-IAA
FRAP/Fixation/Live-cell imaging
0
12 h
13 h
C
1 µg/ml Doxycycline
1 µM 5Ph-IAA + 0.1 µg/ml Nocodazole
Mitotic shake-off
0
12 h
16 h
D
INCENP Survivin Borealin
Sgo1

**Figure S5**

#### Supplementary Figure Legends

**Fig. S1. (A)** Sequence alignment of Borealin orthologues from Homo sapiens (hs), Bos taurus (bt), Mus musculus (mm), Gallus gallus (gg), Danio rerio (dr), and Xenopus laevis (xl). The sequences are colored based on conservation with red being highly conserved and yellow being poorly conserved. The predicted secondary structure elements are shown below the sequence alignment. Multiple sequence alignment was performed with Clustal Omega (EMBL-EBI) and edited with Jalview 2.11.0 (Waterhouse et al., 2009). The secondary structure prediction was done using the web services in Jalview, using Jpred. The N-terminal tail, loop region and dimerization domain of Borealin are highlighted with boxes. **(B)** Sequence alignment of the first 190 amino acids of INCENP orthologues from Homo sapiens (hs), Bos taurus (bt), Mus musculus (mm), Gallus gallus (gg), Danio rerio (dr), and Xenopus laevis (xl). The sequences are colored based on conservation with red being highly conserved and yellow being poorly conserved. The predicted secondary structure elements are shown below the sequence alignment. Multiple sequence alignment was performed with Clustal Omega (EMBL-EBI) and edited with Jalview 2.11.0 (Waterhouse et al., 2009). The secondary structure prediction was done using the web services in Jalview, using Jpred. The basic IDR region of INCENP (including the 'RRKKRR' motif and two additional positive stretches) and the PxVxI motif are highlighted with boxes in the alignment.

**Fig. S2. (A)** SEC chromatogram for $CPC_{I1\text{-}190SB}$-H3T3ph NCP complex formation and the corresponding SDS-PAGE gel for the run. The peak fractions were pooled and used to solve the cryo-EM structure of the complex. **(B)** Representative 2D-classes from the processing pipeline of the cryo-EM structure from CryoSPARC. **(C)** Local resolution map of the cryo-EM density is depicted in a cyan to pink color gradient. The scale bar represents the corresponding resolution in Å. (**D**) GSFSC plot, cFSCs curve, and azimuth plot from CryoSPARC for the cryo-EM density. **(E)** Cryo-EM model surface colored by electrostatic potential (APBS using Pymol version 3.1), with CPC rotated -180 degrees around the y axis to highlight the highly basic Borealin N terminal tail (box, left), which binds to the acidic patch of the nucleosome (box, right).

**Fig. S3. (A)** SDS-PAGE gel of EDC-NHS crosslinked $CPC_{I1\text{-}190SB}$-H3T3ph NCP complex. **(B)** SDS-PAGE gels with the input samples for the UV-crosslinking experiments with H3T3ph

NCPS (top) and 601 Widom 147bp IR700 DNA (bottom). **(C)** Schematic diagram depicting the domain architectures of $CPC_{I1\text{-}190SB}$, $CPC_{I1\text{-}190\_6ASB}$, $CPC_{I1\text{-}190SB\Delta loop}$, $CPC_{I1\text{-}190SB10\text{-}end}$, and $CPC_{I1\text{-}58SB}$. **(D)** Representative SDS-PAGE gel with the $CPC_{I1\text{-}190SB}$, $CPC_{I1\text{-}190\_6ASB}$, and $CPC_{I1\text{-}190SB\Delta loop}$ inputs of EMSAs.

**Fig. S4. (A)** AFM characterization of bare DNA construct and reconstituted H3T3ph nucleosomes. DNA contour length distribution as measured via automated readout. The mode of the distribution (red dashed line) is at contour length 155 nm, corresponding to 0.32 nm/bp, in good agreement with previous AFM measurements of DNA length (Rivetti, et al. JMB 1996; Konrad et al., 2021). The experimental rise per base pair is used to convert the units of wrapped length ΔL from nm to bp. **(B)** Distribution of normalized nucleosome positions, quantified as the ratio of the short arm length $l_1$ over the sum of the arm lengths $l_1$+$l_2$. Only fully wrapped nucleosomes are taken into account because asymmetric nucleosome unwrapping shifts the distribution. The red dashed line indicates the nucleosome position as expected from the DNA construct design. The full black line is a Gaussian fit to the data with a mean value of 0.29. **(C)** Schematic depiction indicating that geometrical convolution by the AFM tip distorts the real structure and leads to an underestimation of the nucleosome opening angle. **(D)** Two-dimensional kernel density estimate with indicated positions of different unwrapping states (numbers of bp unwrapped with respect to the fully wrapped state), as deduced from AFM image simulations (see Konrad et al., Biophys J 2022). **(E)** Native gel with AFM DNA and reconstituted H3T3ph NCPs with AFM DNA. **(F)** SDS-PAGE gel with the inputs for the mono-nucleosome MNase assay in Fig. 5E. **(G)** SDS-PAGE gel with the inputs of the MNase assay with 12-mer nucleosomal arrays in Fig. 5F.

**Fig. S5. (A)** Representative immunoblot for RPE1 cell lines expressing either INCENP wt or INCENP 6A mutant showing the expression levels of the different INCENP constructs used in Fig. 4. **(B)** Diagram with the experimental timeline followed for the RPE1 experiments shown in Fig. 4A, 4C and 4D. **(C)** Diagram with the experimental timeline followed for the RPE1 experiment shown in Fig. 4B. **(D)** CPC-NCP complex model depicted with globular visualization in ChimeraX version: 1.9 (left), and the Alpha fold 3 prediction of SGO motif of Sgo1 ($Sgo1_{466\text{-}527}$) interacting with the H3T120ph nucleosome (right). Based on our model and the alpha fold prediction, CPC binding and Sgo1 binding cannot co-exist due to steric hindrance.
